# Highly Stable Mn(V)-Nitrido and Nitrogen-Atom Transfer Reactivity within a *De Novo* Protein

**DOI:** 10.64898/2026.03.23.713767

**Authors:** Jithin Thomas, Sudha Yadav, Paul H. Oyala, Veronica Carta, David P. Goldberg, Samuel I. Mann

## Abstract

High-valent metal–nitrido species are powerful nitrogen-atom transfer intermediates but remain difficult to access and control due to intrinsic instability and bimolecular N–N coupling pathways. Herein, we report the first formation of a high-valent Mn(V)–nitrido complex within a *de novo* designed protein scaffold and demonstrate that a reactive precursor to this species can be catalytically intercepted for enantioselective aziridination. A Mn(V)≡N unit derived from an abiological diphenyl porphyrin is confined within a designed helical bundle protein, where the protein environment suppresses bimolecular decay and enables detailed spectroscopic characterization. Electron paramagnetic resonance, resonance Raman, and circular dichroism spectroscopies confirm formation of a low-spin Mn(V)–nitrido species that is stable for weeks at room temperature and exhibits minimal perturbation of the Mn≡N unit upon modulation of the axial histidine ligand, while catalytic activity and stereochemical outcome are sensitive to its presence. Mechanistic studies identify monochloramine (NH_2_Cl) as the operative nitrogen-atom donor and support the involvement of a transient Mn-bound N-transfer intermediate en route to nitrido formation. Under catalytic conditions, this intermediate is intercepted to perform aziridination with TON ≈ 180 and an enantiomeric ratio of 65:35. Together, these results establish *de novo* protein design as a platform for stabilizing high-valent metal–nitrido species and harnessing their reactivity for nitrogen-atom transfer chemistry beyond the limits of natural metalloenzymes and small-molecule catalysts.

---

High-valent metal–nitrido species are key intermediates in nitrogen-atom transfer chemistry, yet remain difficult to access and control.^1-11^ Here, we report the first formation of a high-valent Mn(V)-nitrido complex within a *de novo* designed protein scaffold and demonstrate that a reactive precursor to this species - possibly a Mn-nitrenoid - can be intercepted to enable stereochemically biased aziridination under aqueous conditions. Access to metal-nitrido chemistry in biological environments has been historically limited^12-15^ by intrinsic instability, including bimolecular N-N coupling, and by the absence of platforms that allow control over axial ligation, second-sphere interactions, and stereochemical outcomes. While Mn-nitrido porphyrins have previously been generated in native hemoproteins and analyzed by resonance Raman spectroscopy (rR)^16^, such systems did not provide programmable control over metal coordination or enable selective N-atom transfer to an external substrate. The present work establishes *de novo* proteins as a platform to both stabilize high-valent nitrido intermediates and harness their reactivity in ways that remain challenging for small-molecule and biocatalytic systems (**Fig. 1**).

**Figure 1.**
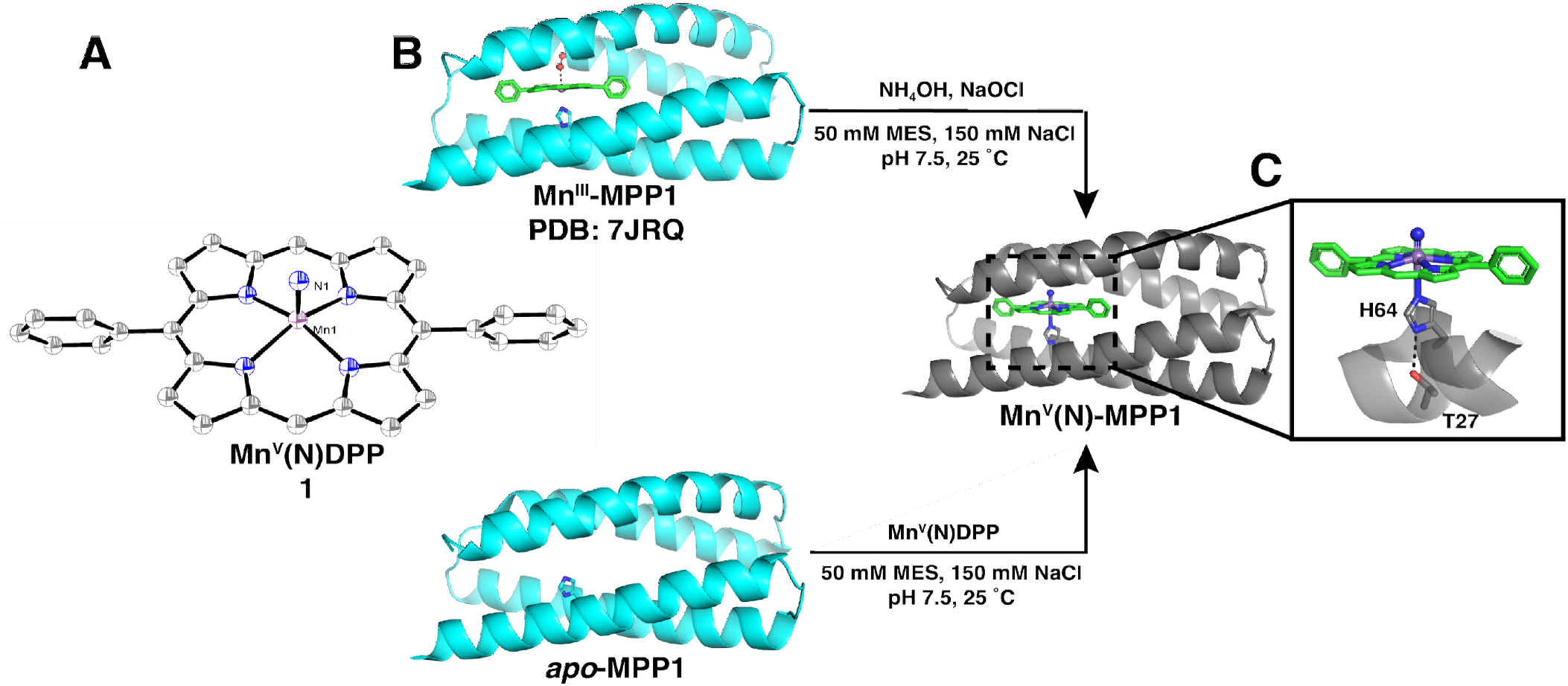
(A) ORTEP diagram (at 50% probability level) of Mn(V)DPP (**1**) (H-atoms have been omitted for clarity). (B) Scheme showing two methods to generate Mn(V)N-MPP1: (top) direct oxidation of Mn(III)-MPP1 and (bottom) addition of pre-formed **1** to apo-MPP1. (C) PyMol model of **1** bound within MPP1 constructed from PDB: 7JRQ.

*De novo* protein design provides a powerful strategy for stabilizing and interrogating reactive metal intermediates within precisely defined environments that are inaccessible to both natural enzymes and purely synthetic ligands. Previously, we showed that a designed helical bundle protein (MPP1) can accommodate an abiological porphyrin cofactor, manganese diphenyl porphyrin (Mn(III)DPP), and support transient formation of a high-valent Mn(V)-oxo species in approximately 60% yield, demonstrating that *de novo* protein scaffolds can be engineered to control high-valent metal chemistry.^17-18^ In this report, we apply this design paradigm to a chemically distinct class of high-valent intermediates by preparing and stabilizing, to our knowledge for the first time, a Mn(V)-nitrido complex within a *de novo* protein. This advance enables direct spectroscopic interrogation of metal-nitrogen multiple bonding under programmable axial ligation and confinement and illustrates how protein environments can be leveraged to stabilize and exploit metal-nitrenoid reactivity. *De novo* protein scaffolds allow axial ligands to be positioned in close proximity to the metal center, enabling conditional interactions that can be engaged or disengaged along a catalytic reaction coordinate. Notably, most small-molecule and biocatalytic N-atom transfer systems rely on pre-activated N-alkyl (NR) nitrene equivalents, often requiring strong acids.^6, 8, 15, 19-24^ By contrast, the present system effects N–H transfer under aqueous conditions utilizing aqueous NH_3_,^11^ highlighting a distinct and biologically relevant regime of metal–nitrogen chemistry accessible within a designed protein environment.

We first examined formation of a Mn(V)-nitrido species using the less sterically encumbered Mn(III)DPP cofactor in the absence of a protein, analogous to prior studies on tetraphenyl porphyrin systems.^4-5^ Treatment of Mn(III)DPP with excess NaOCl and NH_4_OH^11^ in dichloromethane results in rapid spectral changes, including a shift of the Soret band from λ_max_ = 458 to 407 nm and loss of features at λ_max_ = 550 nm, consistent with formation of a high-valent Mn-nitrido species (**Figure S1**).^4, 25^

Data obtained from single-crystal X-ray diffraction methods confirmed the structure of Mn(V)N-DPP (**Figure 1A**), revealing a five-coordinate manganese center with average Mn-N_por_ bond lengths of 2.02 Å and a short Mn-N_nitrido_ distance of 1.54 Å, consistent with the assignment of a Mn≡N triple bond (Table S1-S2).^25^ The Mn ion is displaced 0.38 Å out of the porphyrin N4 plane (towards the nitrido ligand) and the Mn1 and N1 are disordered approximately 50:50 above and below the porphyrin plane, indicative of minimal steric bias in the absence of a structured environment (see SI). In DMSO solution, Mn(V)N-DPP decays over two days to a mixture of Mn(III) and Mn(V) species (**Figure S2**), consistent with bimolecular decomposition pathways.^2, 26^

We next investigated the effects of confining **1** within the *de novo* designed protein scaffold. MPP1 binds Mn(III)DPP in a 1:1 stoichiometry via axial histidine coordination^17^ (**Figure 1B**) and subsequent reaction of Mn(III)-MPP1 with excess NaOCl and NH_4_OH in MES buffer led to formation of a new species exhibiting absorption features at λ_max_ = 406, 530 and 565 nm (**Figure 2A**), closely matching those observed for Mn(V)N-DPP in solution (**Figure S1**). This species was therefore assigned as Mn(V)N-MPP1.

**Figure 2.**
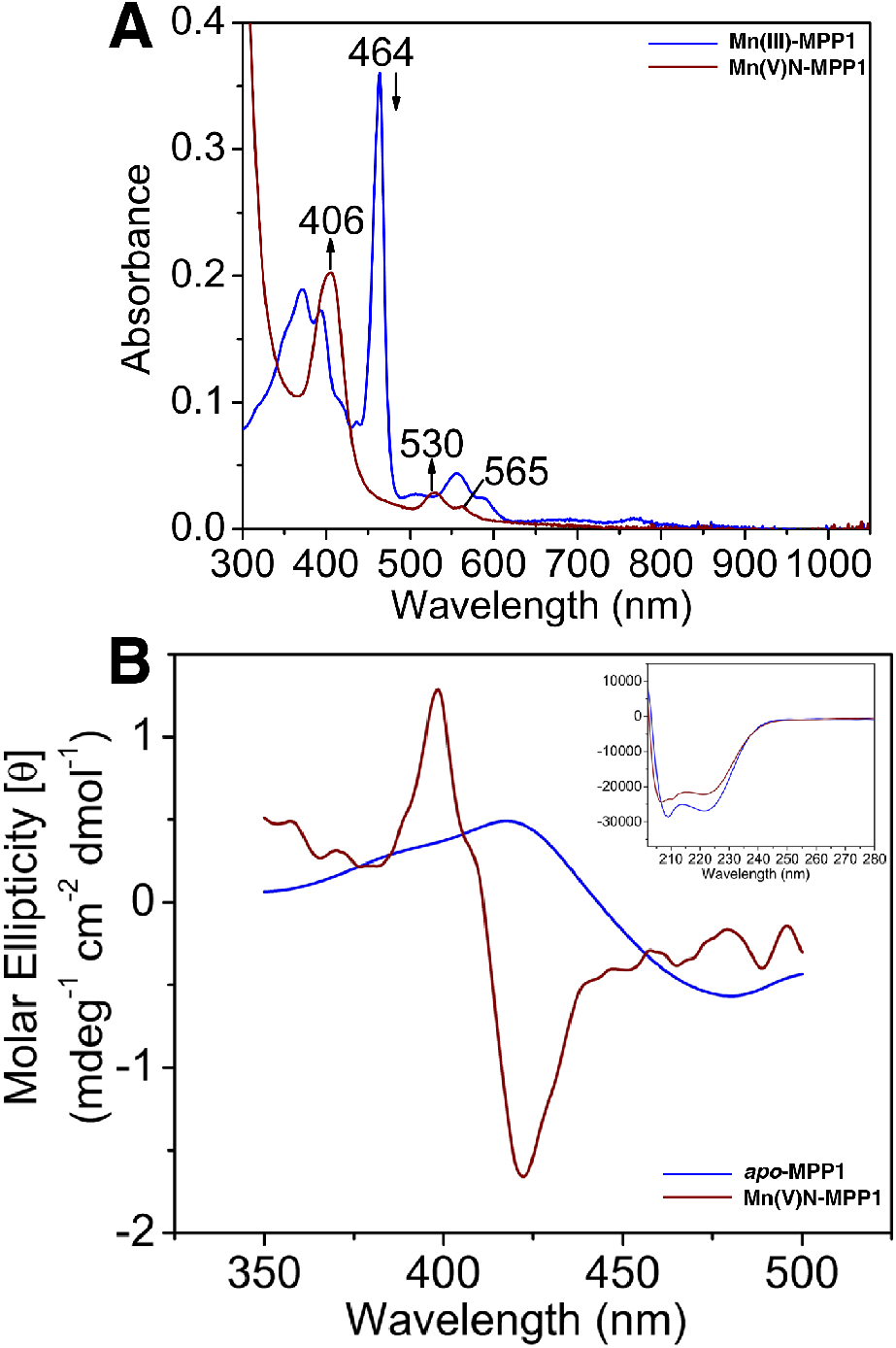
(a) Electronic absorption spectra of Mn(III)MPP1(blue) and Mn(V)N-MPP1 (red). (b) Visible-CD spectra of Mn(V)N-MPP1 and apo-MPP1 (inset, UV-CD spectrum of Mn(V)N-MPP1 and apo-MPP1).

Independent preparation of Mn(V)N-MPP1 by incubation of pre-formed solution of crystalline Mn(V)N-DPP in DMSO with apo-MPP1 yielded an identical absorption spectrum (**Figure S3**), further indicating that the protein scaffold supports binding of the nitrido complex. In contrast to the free cofactor, Mn(V)N-MPP1 is stable at room temperature for at least 3 months with no evidence of bimolecular decay (**Figure S4**), demonstrating that confinement within the designed protein effectively suppresses decomposition pathways.

Although Mn(V)N-DPP is achiral in solution, binding within the protein active site is expected to induce chirality if the cofactor is held in a rigid orientation. A structure-guided model of Mn(V)N-MPP1 created by an overlay of the structure of MnNDPP with the MnDPP within MPP1 (PDB: 7JRQ, **Figure 1C** and **4C**) predicts the nitrido ligand sits within a rigid, non-polar environment with a trans-axial His ligand. Indeed, the visible circular dichroism spectrum of Mn(V)N-MPP1 exhibits a pronounced Cotton effect associated with the Soret transition at λ_max_ = 406 nm, coincident with the electronic absorption features. Far-UV CD measurements confirm that the secondary structure of MPP1 remains intact following nitrido formation (**Figure 2B**), indicating that the protein scaffold tolerates the strongly basic conditions required for Mn(V)–nitrido generation.

Formation of the Mn(V)-nitrido species in MPP1 is accompanied by a loss of the six-line Mn(III) X-band parallel-mode EPR signal centered at g = 2.011 observed for the resting Mn(III)-MPP1 complex (**Figure S5**).^27^ No signal is observed for Mn(V)N-MPP1 in either parallel- or perpendicular-mode X-band EPR, consistent with a diamagnetic, low-spin *d*^2^ electronic configuration (**Figure S6**).^28^ These observations further support assignment of a Mn(V) oxidation state and a closed-shell Mn≡N unit.

Positioning of an axial histidine ligand in MPP1 provides a rare opportunity to directly probe axial ligation effects on a metal-nitrido unit within a well-defined environment. The rR spectra of Mn(V)N-MPP1 reveal Mn-N stretching modes at 1049 cm^-1^ (**Figure 3**), closely matching those observed for the free Mn(V)N-DPP complex (1048 cm^-1^, **Figure S7**) which we attribute to the hydrophobic environment of MPP1.^16, 29^ Isotopic labeling using ^15^NH_4_Cl resulted in a down shift of 28 cm^-1^ (calculated shift 28 cm^-1^), confirming assignment of these modes to the Mn≡N stretch. Despite the proximity of the axial histidine residue suggested by the protein structure (**Figure 1C** and **4C**), these data indicate that the axial histidine has a minimal effect on the Mn≡N bond strength. To further test this hypothesis, we prepared an H64A variant of MPP1 lacking the axial histidine ligand. The absorption spectrum for Mn(III)MPP1-H64A shows shifted Soret and Q-band absorbances, consistent with removal of the axial His ligand (**Figure S9**). However, formation of Mn(V)N-MPP1-H64A yields absorption and rR spectra that match those of the wild-type protein (**Figure S8**-**S9**), confirming that H64 does not significantly perturb the Mn≡N unit. We hypothesize that the displacement of the Mn-center out of the porphyrin N4 plane results in weak or no interaction with the His residue. In MPP1 (PDB: 7JRQ) the Mn is displaced 0.14 Å towards the axial His (**Figure 4A**), while in MnDPP the Mn is displaced 0.38 Å towards the nitrido ligand (**Figure 1C/4C**). These results highlight the intrinsic robustness of the nitrido bond and underscores the utility of *de novo* protein design for systematically interrogating axial ligand effects that are difficulty to access in synthetic systems.^30^

**Figure 3.**
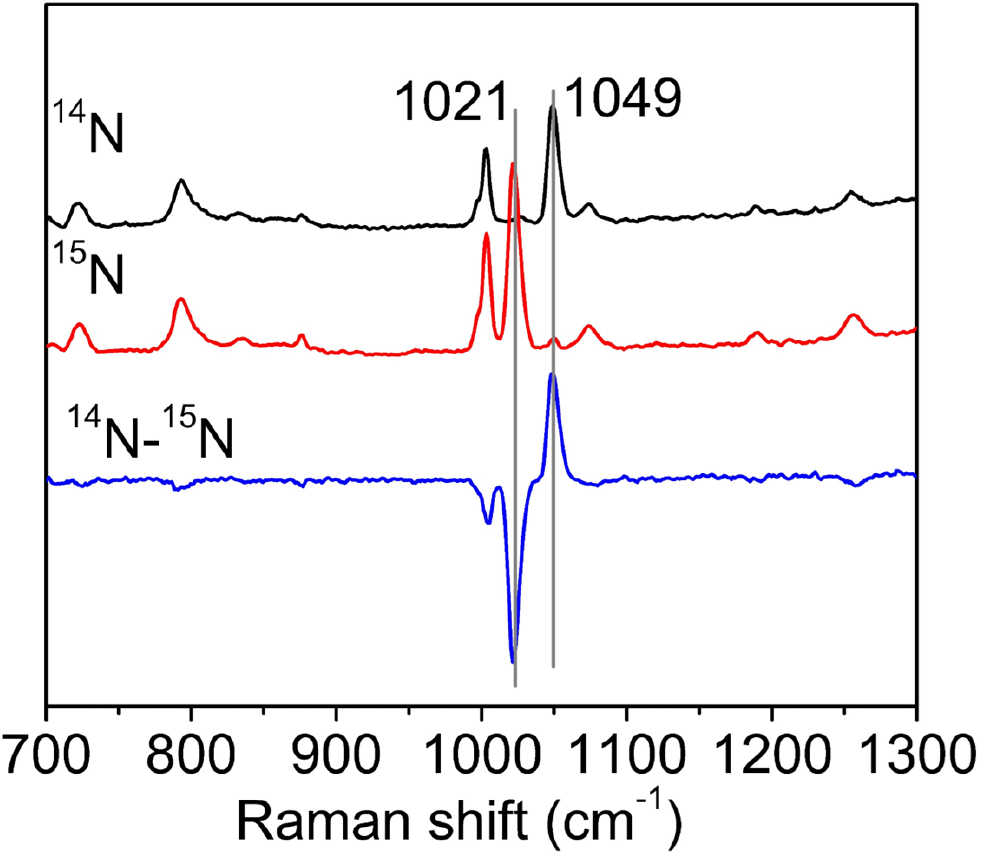
rR data for Mn(V)N MPP1(WT) showing ^14^N (black), ^15^N (red), and difference spectrum (blue).

**Figure 4.**
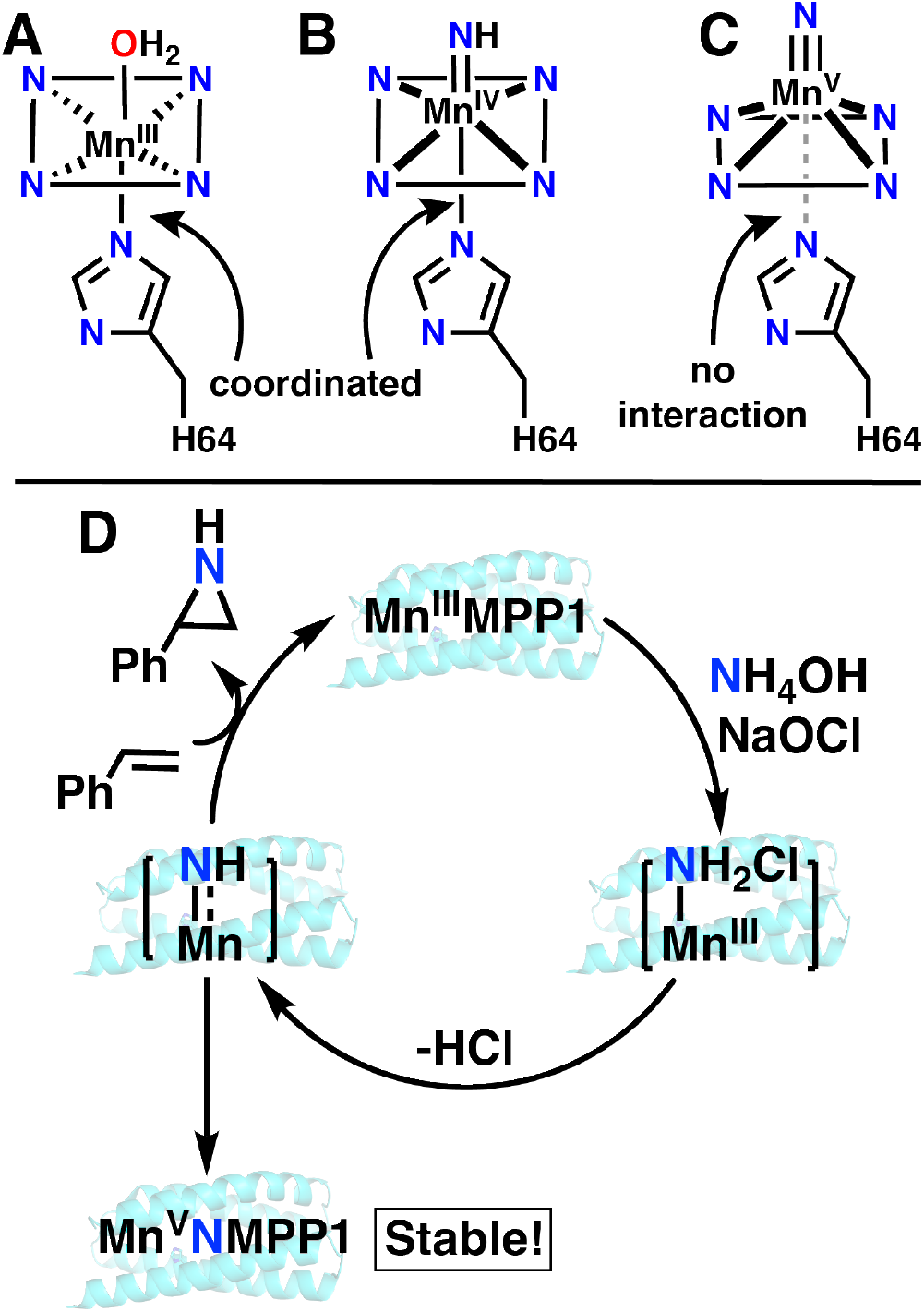
Illustration of proposed role of His. (A) In the resting state Mn(III)-aqua state, H64 serves as an axial ligand as observed crystallographically. (B) During catalysis, a transient, unobserved Mn-nitrene intermediate may access axial coordination due to reduced metal displacement above the porphyrin plane. (C) In the thermodynamically stable Mn(V)N state, strong metal-nitrogen multiple bonding displaces the Mn center toward the nitrido ligand, resulting in minimal interaction with H64. (D) Proposed catalytic cycle for aziridination.

We next sought to identify the nitrogen-atom donor responsible for formation of the Mn(V)-nitrido species under the NaOCl/NH_4_OH conditions. Mixing hypochlorite and ammonia is known to generate monochloramine (NH_2_Cl), a highly reactive and transient gaseous species that has been implicated in nitrene and nitrido formation in synthetic systems.^31-35^ We therefore hypothesized that NH_2_Cl is the operative reagent responsible for the Mn-N bond formation in Mn(V)N-MPP1. To directly test this hypothesis, NH_2_Cl was generated *in situ* and trapped by bubbling the evolved gas into diethyl ether at -80 °C, affording an ethereal solution of NH_2_Cl.^31^ Addition of one equivalent of NH_2_Cl to Mn(III)-MPP1 resulted in rapid formation of Mn(V)N-MPP1, as judged by the appearance of characteristic absorption bands at λ_max_ = 407 and 530 nm that are indistinguishable from those obtained by using NaOCl/NH_4_OH (**Figure S10a**). These results strongly support NH_2_Cl as the nitrogen-atom transfer reagent responsible for nitrido formation within the protein scaffold. Formation of Mn(V)-nitrido from NH_2_Cl implies the intermediacy of a more reactive nitrogen-containing species, potentially a Mn-nitrene^36-37^, en route to the thermodynamically stable nitrido complex. We therefore examined whether this intermediate could be intercepted for catalysis prior to the Mn≡N formation.

While the isolated Mn(V)N-MPP1 complex is unreactive toward styrene (**Scheme 1, Figure S11**), reaction of Mn(III)DPP-MPP1 with NaOCl/NH_4_OH (or NH_2_Cl) in the presence of excess styrene for 5 minutes results in catalytic aziridination accompanied by suppression of Mn(V)-N accumulation (**Scheme 1, Figure S10, S12-S13**). Under these conditions (25 µM Mn-MPP1, 10 mM styrene), aziridine-derived products are formed within 5 min in ∼50% yield relative to styrene, corresponding to approximately 180 turnovers per Mn center (**Table S3 and Figure S14**). UPC and GC-MS analysis identify N-chloro-2-phenylaziridine as the major nitrogen-containing product, arising from rapid N-chlorination of the initially formed aziridine by excess hypochlorite under the reaction conditions (**see SI**). This product is formed with an enantiomeric ratio (*er*) of R:S = 65:35, indicating that the protein environment imparts stereo-chemical bias during aziridination (**Fig. S15**). Control experiments demonstrate that racemic 2-phenylaziridine undergoes non-enantioselective N-chlorination with NaOCl, confirming that the measured *er* reflects the stereochemical outcome of the protein-mediated aziridination step (**Fig. S16**). Formation of (1,2-dichloroethyl)benzene is also observed and arises from the background reaction of styrene with NaOCl in the absence of protein (**Figure S14**). These results demonstrate that a transient Mn-bound N-transfer intermediate generated prior to Mn(V)-nitrido formation can be catalytically intercepted within the chiral environment of the *de novo* protein scaffold to effect modest enantioselective aziridination under aqueous conditions.

**Scheme 1.**
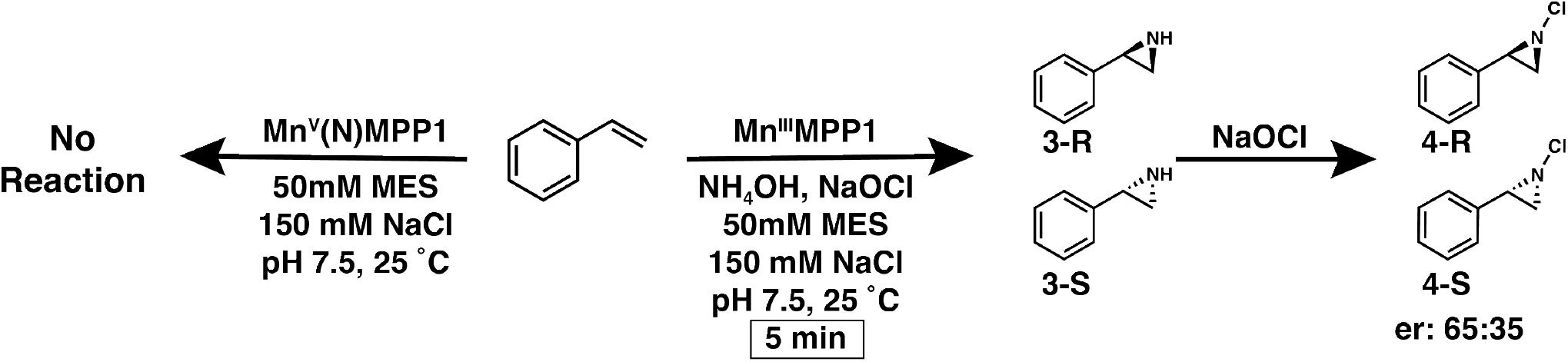
Aziridination of Styrene by MnMPP1 and Mn(V)NMPP1

Notably, the H64A variant of MPP1, despite showing no effect on the Mn(V)-nitrido spectroscopic properties relative to WT, shows substantially diminished aziridination activity under identical conditions (∼15% yield, **Table S3)**. This decoupling of Mn≡N bond perturbation from catalytic performance suggests that H64 is not a primary determinant of nitrido bond strength but instead may influence the formation, lifetime, or productive use of the transient N-transfer intermediate that precedes Mn(V)-nitrido formation (**Figure 4B**). The proximity of H64 within the designed scaffold enables designed, conditional axial interactions that may become accessible along the reaction coordinate, when the manganese center is expected to reside closer to the porphyrin plane (**Figure 4B**). In this regime, axial coordination could provide a level of dynamic regulation that is difficult to achieve in purely synthetic systems.

In summary, we report the first example of a high-valent metal-nitrido species stabilized within a *de novo* protein scaffold. Confinement of a Mn(V)-nitrido porphyrin within MPP1 suppresses bimolecular decomposition pathways observed for the free cofactor and enables comprehensive spectroscopic characterization of a Mn≡N unit under protein-defined constraints. Interrogation of the axial ligation using wild-type and H64A variants reveals that axial histidine coordination has minimal impact on the nitrido bond strength, highlighting the intrinsic robustness of the Mn≡N multiple bond and demonstrating the level of ligand and secondary-structure control that is difficult to access in purely synthetic systems. Mechanistic studies identify NH_2_Cl as the operative nitrogen-atom donor responsible for nitrido formation and provide evidence for a transient, high-energy N-transfer intermediate that precedes Mn(V)-nitrido formation. Importantly, this intermediate can be utilized within the chiral protein environment to impart stereo-chemical bias during aziridine formation (**Fig. 4D**). More broadly, this work highlights a design principle in which axial ligands can be positioned in proximity to a metal center without enforcing persistent coordination, enabling dynamic engagement during catalysis while preserving the stability of high-valent intermediates. Together, these results establish *de novo* protein design as a powerful platform for accessing, stabilizing, and mechanistically interrogating high-valent metal-nitrido species, opening new opportunities to explore nitrogen-transfer chemistry beyond the limits of natural metalloenzymes.

## ASSOCIATED CONTENT

### Supporting Information

The Supporting Information is available free of charge on the ACS Publications website.

Materials, experimental methods, and spectroscopic characterization data including Figures S1−S38 and Tables S1−S6.

CCDC 2528049 contains the supplementary crystallographic data for this paper. These data can be obtained free of charge via www.ccdc.cam.ac.uk/data_request/cif, or by emailing, or by contacting The Cambridge Crystallographic Data Centre, 12 Union Road, Cambridge CB2 1EZ, UK; fax: +44 1223 336033.

## ACKNOWLEDGMENT

We thank Prof. Ana Bahamonde and Lang Cheng (Oscar) Hung for access to and help running their UPC.

---

Insert Table of Contents artwork here

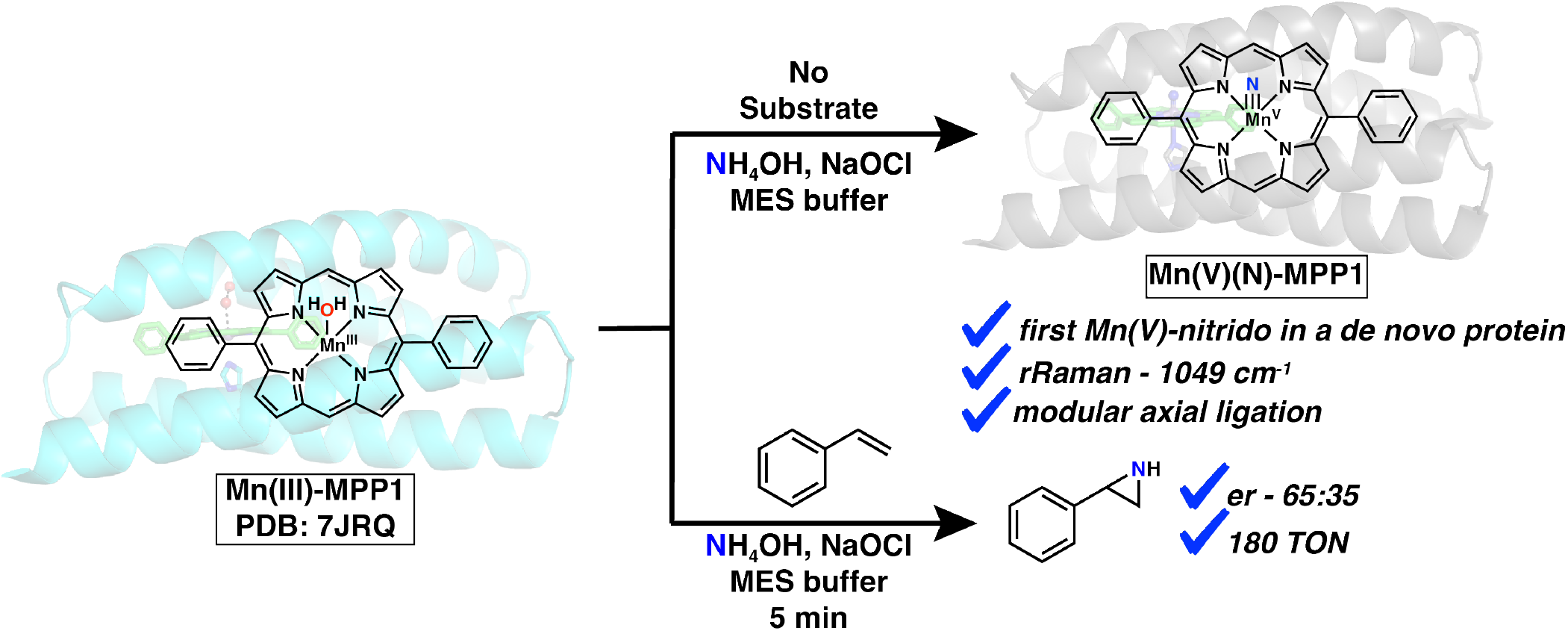

## REFERENCES

1. Eikey Rebecca, A.; Khan, S. I.; Abu-Omar Mahdi, M., The elusive terminal imido of manganese(V). Angew. Chem. Int. Ed. 2002, 41, 3592–3595.

2. Xiang, J.; Shi, H.; Man, W. L.; Lau, T. C., Design of Highly Electrophilic and Stable Metal Nitrido Complexes. Acc. Chem. Res. 2024, 57, 2700–2716.

3. Cosio, M. N.; Powers, D. C., Prospects and challenges for nitrogen-atom transfer catalysis. Nat. Rev. Chem. 2023, 7, 424–438.

4. Hill, C. L.; Hollander, F. J., Structural characterization of nitrido[tetrakis(p-methoxyphenyl)porphinato]manganese(V). J. Am. Chem. Soc. 1982, 104, 7318–7319.

5. Du Bois, J.; Tomooka, C. S.; Hong, J.; Carreira, E. M., Nitridomanganese(V) Complexes: Design, Preparation, and Use as Nitrogen Atom-Transfer Reagents. Acc. Chem. Res. 1997, 30, 364–372.

6. Goswami, M.; Lyaskovskyy, V.; Domingos, S. R.; Buma, W. J.; Woutersen, S.; Troeppner, O.; Ivanovic-Burmazovic, I.; Lu, H.; Cui, X.; Zhang, X. P.; Reijerse, E. J.; DeBeer, S.; van Schooneveld, M. M.; Pfaff, F. F.; Ray, K.; de Bruin, B., Characterization of Porphyrin-Co(III)-’Nitrene Radical’; Species Relevant in Catalytic Nitrene Transfer Reactions. J. Am. Chem. Soc. 2015, 137, 5468–5479.

7. Leung, S. K.; Huang, J. S.; Liang, J. L.; Che, C. M.; Zhou, Z. Y., Nitrido ruthenium porphyrins: synthesis, characterization, and amination reactions with hydrocarbon or silyl enol ethers. Angew. Chem. Int. Ed.Engl. 2003, 42, 340–343.

8. Roizen, J. L.; Harvey, M. E.; Du Bois, J., Metal-catalyzed nitrogen-atom transfer methods for the oxidation of aliphatic C-H bonds. Acc. Chem. Res. 2012, 45, 911–922.

9. Leeladee, P.; Goldberg, D. P., Epoxidations catalyzed by manganese(V) oxo and imido complexes: role of the oxidant-Mn-oxo (imido) intermediate. Inorganic Chemistry 2010, 49, 3083–3085.

10. Lansky, D. E.; Kosack, J. R.; Narducci Sarjeant, A. A.; Goldberg, D. P., An Isolable, Nonreducible High-Valent Manganese(V) Imido Corrolazine Complex. Inorg. Chem. 2006, 45, 8477–8479.

11. Shi, H.; Lee, H. K.; Pan, Y.; Lau, K. C.; Yiu, S. M.; Lam, W. W. Y.; Man, W. L.; Lau, T. C., Structure and Reactivity of a Manganese(VI) Nitrido Complex Bearing a Tetraamido Macrocyclic Ligand. J. Am. Chem. Soc. 2021, 143, 15863–15872.

12. Svastits, E. W.; Dawson, J. H.; Breslow, R.; Gellman, S. H., Functionalized nitrogen atom transfer catalyzed by cytochrome P-450. J. Am. Chem. Soc. 1985, 107, 6427–6428.

13. Coin, G.; Latour, J. M., Nitrene transfers mediated by natural and artificial iron enzymes. J. Inorg. Biochem. 2021, 225, 111613.

14. Brandenberg, O. F.; Fasan, R.; Arnold, F. H., Exploiting and engineering hemoproteins for abiological carbene and nitrene transfer reactions. Curr. Opin. Biotechnol. 2017, 47, 102–111.

15. Yang, Y.; Arnold, F. H., Navigating the Unnatural Reaction Space: Directed Evolution of Heme Proteins for Selective Carbene and Nitrene Transfer. Acc. Chem. Res. 2021, 54, 1209–1225.

16. Tsubaki, M.; Hori, H.; Hotta, T.; Hiwatashi, A.; Ichikawa, Y.; Yu, N. T., Influence of heme-surrounding amino acid residues on the manganese (V)-nitrido bond in manganese-substituted hemoproteins: resonance Raman evidence for porphyrin core expansion and reduction of the manganese(V)-nitrido stretching force constant. Biochemistry 1987, 26, 4980–4986.

17. Mann, S. I.; Nayak, A.; Gassner, G. T.; Therien, M. J.; DeGrado, W. F., De Novo Design, Solution Characterization, and Crystallographic Structure of an Abiological Mn-Porphyrin-Binding Protein Capable of Stabilizing a Mn(V) Species. J. Am. Chem. Soc. 2021, 143, 252–259.

18. Coronado, K. R.; Zhu, Y.; Mann, S. I., De novo design of four-helix bundle proteins to bind metalloporphyrin cofactors. Methods Enzymol. 2025, 720, 1–22.

19. Farwell, C. C.; Zhang, R. K.; McIntosh, J. A.; Hyster, T. K.; Arnold, F. H., Enantioselective Enzyme-Catalyzed Aziridination Enabled by Active-Site Evolution of a Cytochrome P450. ACS Cent. Sci. 2015, 1, 89–93.

20. Athavale, S. V.; Gao, S.; Das, A.; Mallojjala, S. C.; Alfonzo, E.; Long, Y.; Hirschi, J. S.; Arnold, F. H., Enzymatic Nitrogen Insertion into Unactivated C-H Bonds. J. Am. Chem. Soc. 2022, 144, 19097–19105.

21. Gao, S.; Das, A.; Alfonzo, E.; Sicinski, K. M.; Rieger, D.; Arnold, F. H., Enzymatic Nitrogen Incorporation Using Hydroxylamine. J. Am. Chem. Soc. 2023, 145, 20196–20201.

22. Yang, Y.; Arnold, F. H., Navigating the Unnatural Reaction Space: Directed Evolution of Heme Proteins for Selective Carbene and Nitrene Transfer. Acc. Chem. Res. 2021, 54, 1209–1225.

23. Hyster, T. K.; Farwell, C. C.; Buller, A. R.; McIntosh, J. A.; Arnold, F. H., Enzyme-controlled nitrogen-atom transfer enables regiodivergent C-H amination. J. Am. Chem. Soc. 2014, 136, 15505–15508.

24. Goldberg, N. W.; Knight, A. M.; Zhang, R. K.; Arnold, F. H., Nitrene Transfer Catalyzed by a Non-Heme Iron Enzyme and Enhanced by Non-Native Small-Molecule Ligands. J. Am. Chem. Soc. 2019, 141, 19585–19588.

25. Groves, J. T.; Takahashi, T.; Butler, W. M., Synthesis and molecular structure of a nitrido(porphyrinato)chromium(V) complex. Inorg. Chem. 1983, 22, 884–887.

26. Seymore, S. B.; Brown, S. N., Polar effects in nitride coupling reactions. Inorg. Chem. 2002, 41, 462–469.

27. Campbell, K. A.; Yikilmaz, E.; Grant, C. V.; Gregor, W.; Miller, A.-F.; Britt, R. D., Parallel Polarization EPR Characterization of the Mn(III) Center of Oxidized Manganese Superoxide Dismutase. J. Am. Chem. Soc. 1999, 121, 4714–4715.

28. Groves, J. T.; Takahashi, T., Activation and transfer of nitrogen from a nitridomanganese(V) porphyrin complex. Aza analog of epoxidation. J. Am. Chem. Soc. 1983, 105, 2073–2074.

29. Campochiaro, C.; Hofmann, J. A.; Bocian, D. F., Resonance Raman spectra of chromium(V) and manganese(V) porphyrin nitrides. Inorg. Chem. 1985, 24, 449–450.

30. Thomas, J.; Mokkawes, T.; Senft, L.; Dey, A.; Gordon, J. B.; Ivanovic-Burmazovic, I.; de Visser, S. P.; Goldberg, D. P., Axial Ligation Impedes Proton-Coupled Electron-Transfer Reactivity of a Synthetic Compound-I Analogue. J. Am. Chem. Soc. 2024, 146, 12338–12354.

31. Hynes, J., Jr.; Doubleday, W. W.; Dyckman, A. J.; Godfrey, J. D., Jr.; Grosso, J. A.; Kiau, S.; Leftheris, K., N-Amination of pyrrole and indole heterocycles with monochloramine (NH2Cl). J. Org. Chem. 2004, 69, 1368–1371.

32. Meyer, K.; Bendix, J.; Metzler-Nolte, N.; Weyhermueller, T.; Wieghardt, K., Nitridomanganese(V) and -(VI) Complexes Containing Macrocyclic Amine Ligands. J. Am. Chem. Soc. 1998, 120, 7260–7270.

33. Qiang, Z.; Adams, C. D., Determination of monochloramine formation rate constants with stopped-flow spectrophotometry. Environ. Sci. Technol. 2004, 38, 1435–1444.

34. Trofe, T. W.; Inman, G. W.; Johnson, J. D., Kinetics of monochloramine decomposition in the presence of bromide. Environ. Sci. Technol. 2002, 14, 544–549.

35. Weil, I.; Morris, J. C., Kinetic Studies on the Chloramines. The Rates of Formation of Monochloramine, N-Chlormethylamine and N-Chlordimethylamine. J. Am. Chem. Soc. 1949, 71, 1664–1671.

36. Das, A.; Gao, S.; Lal, R. G.; Hicks, M. H.; Oyala, P. H.; Arnold, F. H., Reaction Discovery Using Spectroscopic Insights from an Enzymatic C-H Amination Intermediate. J. Am. Chem. Soc. 2024, 146, 20556–20562.

37. Van Trieste, G. P., 3rd; Reid, K. A.; Hicks, M. H.; Das, A.; Figgins, M. T.; Bhuvanesh, N.; Ozarowski, A.; Telser, J.; Powers, D. C., Nitrene Photochemistry of Manganese N-Haloamides*. Angew. Chem. Int. Ed. Engl. 2021, 60, 26647–26655.

