## Supporting Information for "Highly Stable Mn(V)-Nitrido and Nitrogen-Atom Transfer Reactivity within a *De Novo* Protein"

**Table of Contents**


1. Experimental Section
2. Spectroscopic Characterization
   - UV-vis studies
   - EPR Analysis
   - Circular Dichroism Analysis
   - rRaman studies
   - X-Ray structure
3. Reactivity and Catalytic Studies
   - Control experiments
   - Reduction experiments
   - pH experiments
   - Calibration curves of standard products
   - Enantioselectivity and absolute configuration of products determination
   - Site directed mutagenesis H64A studies
   - Stoichiometric experiments using *in-situ* generated $NH_2Cl$

### Experimental Section

#### General Methods

All commercial reagents and solvents were obtained from Sigma Aldrich and other commercial providers and unless otherwise indicated used as received. $NH_2Cl$ was prepared using a reported procedure and stored at -80 °C in diethyl ether.[4] The $^{15}NH_4Cl$ was obtained from ICON Cambridge isotopes, Amicon® and Centricon® filters were obtained from EMD Millipore and Mn(III) diphenyl porphyrin chloride was obtained from Frontier Scientific. Silica gel chromatography was carried out using Silia Flash Irregular Silica Gel P60, 40 - 63 µm, 60 Å (R12030B). MPP1 was expressed and purified as previously reported.[6]

#### Instrumentation

UV-visible spectroscopy was carried out with a Cary 60 UV-vis spectrophotometer using a 1 cm modified Schlenk cuvette. Electron paramagnetic resonance (EPR) spectra were acquired using a Bruker EMX CW-EPR spectrometer at a temperature of 5 K. Temperature control was achieved with a continuous-flow liquid helium cryostat (ESR900) and ITC503 temperature controller made by Oxford Instruments, Inc. Resonance Raman spectra were obtained using a custom I302C 500mW ML-UV Kr laser (RoHS) equipped with a liquid-nitrogen-cooled CCD detector (LN-1100 PB, Princeton Instrument). RR spectra were recorded using 406.9 nm excitation in a 180$^{o}$ backscattering geometry on samples maintained at 110 K inside a copper cold finger cooled with liquid nitrogen. A long-pass filter was placed in front of the spectrograph entrance slit to attenuate the Rayleigh scattering. $^{1}H$ NMR spectra were recorded on a Bruker 600 MHz nuclear magnetic resonance spectrometer in $CDCl_3$.GC-MS experiments were performed using Agilent 5975 MSD with Agilent 7890A GC instrument.

#### Preparation of cofactor Mn(V)(N)(DPP), MnMPP1, Mn(V)(N) MPP1

##### a) Synthesis and crystallization of Mn(V)($^{14}$N)(DPP)

The manganese(V) nitrido, 5,15-diphenyl porphyrin, Mn(V)(N)(DPP), was prepared by using a modified version of previously reported procedures for metal nitrido complexes in the literature.[7-8] Mn(Cl)(DPP) (17.5 mg, 2.65 mmol) was dissolved in methanol (3 ml) and eluted down an alumina column with methanol. The methanol was removed and the product was redissolved in 50 ml dichloromethane. This solution was treated with 21 ml of ammonia solution made by diluting 1 ml of concentrated ammonia (14.8 M) with 20 ml of water and allowed to stir for fifteen minutes. A 15 % sodium hypochlorite solution (3 ml) was added, and the reaction was stirred an additional 15 minutes, resulting in a purple solution. The solution was then extracted with two 100 ml portions of water to remove the excess ammonia and hypochlorite and the sodium chloride formed during the reaction. The organic layer was placed on a neutral alumina column, and the product was eluted with dichloromethane. Unreacted manganese (III) porphyrin can be recovered by eluting with methanol. The product was dried under reduced pressure. Crystals suitable for diffraction were obtained by layering chlorobenzene with heptane for 2 days. (Molar extinction coefficient in DCM

= $5.1 \times 10^4$ $M^{-1}$ $cm^{-1}$). $^1H$ NMR (600 MHz, $CDCl_3$) δ 10.21-10.09 (s, 2H, meso-H), 9.45 - 9.08 (s, 8H, β-pyrrole H), 8.30 - 8.02 (m, 4H, o-ArH), 7.95 -7.38 (m, 6H, m/p-ArH), ESI-MS ($+^{ve}$ mode, in DCM/MeOH): *m/z* observed are 515.1(MnDPP) and 530.1[Mn(N)DPP + $H^+$].

**b) Synthesis and crystallization of Mn(V)($^{15}$N)(DPP)**

Mn(Cl)(DPP) (9.3 mg) was dissolved in methanol and eluted down an alumina column with methanol. The methanol was removed and the product was redissolved in 10-15 ml dichloromethane. $^{15}NH_4Cl$ (360 mg) was dissolved in 5 ml ice cold water containing NaOH (550 mg). This solution was mixed for 5 minutes at 0 ˚C and transferred to the round bottom containing Mn(Cl)(DPP). A 15 % sodium hypochlorite solution (5 ml) was added, and the reaction was stirred an additional 15 minutes, resulting in a purple solution. The solution was then washed with two 100 ml portions of water to remove the excess ammonia, hypochlorite, and sodium chloride formed during the reaction. The organic layer was placed on a neutral alumina column, and the product was eluted with dichloromethane.

**c) Preparation of Mn(III)MPP1**

Mn(III)MPP1 was prepared according to previously reported procedures[6] by incubating apo-MPP1 (225 μM, MES buffer, pH 7.5) with Mn(III) diphenylporphyrin chloride (1.1 equiv, 10 mM stock concentration in DMSO solvent). After incubation for 5 minutes, the red protein solution was centrifuged at 14,000 rpm for 3 minutes to remove any residual cofactor and spin filtered to remove the DMSO solvent. This Mn(III)MPP1 was then stored in 4 °C fridge until further use.

**d) Preparation of Mn(V)(N) MPP1**

**Method 1**: Mn(III)MPP1 (18 μM, 10 mL MES buffer) was mixed with $NH_4OH$ (740 mM, 50 μL) and NaOCl (475 mM, 250 μL) at room temperature. The solution was mixed for 5 minutes upon which formation of bubbles, indicative of gaseous $NH_2Cl$ formation, was observed. The color of the solution changed from red to purple during this time. The purple-colored solution was then centrifuged at 14,000 rpm for 3 minutes and washed via spin filter using MES buffer, pH 7.5 for 5-6 times. The Mn(V)NMPP1 was then stored at 4 °C fridge until further use.

**Method 2**: Mn(V)N MPP1 can also be prepared by incubating the apo-MPP1 (40 μM in 1mL MES buffer, pH 7.5) with Mn(V)(N)DPP cofactor (1.2 equiv, 0.36 mM stock concentration in DMSO). After incubation for 15 minutes, the purple protein solution was centrifuged at 14000 rpm for 3 minutes to remove any residual cofactor and washed via spin filter to remove the DMSO.

Note: It is important to prepare stock solutions of Mn(V)(N) DPP in DMSO immediately prior to use. Mn(V)(N)(DPP) was found to partially decompose upon prolonged dissolution (1-2 days) in DMSO, resulting in the formation of a mixture of Mn(III) and Mn(V) species.

**Preparation of EPR Samples**

Mn(III)MPP1 and Mn(V)(N)(MPP1) solution, prepared as described above, were concentrated to a volume of 300 µL using Amicon® spin filters to a concentration of 1-2 mM in 50 mM MES, 150 mM NaCl pH 7.5. An aliquot of the protein solution was taken for UV-vis analysis to confirm Mn(III)MPP1 and formation of the Mn(V)(N). The protein solution was added to an EPR tube and slowly frozen in liquid $N_2$.

**Circular dichroism (CD)**. CD spectra were collected on a Jasco J-815 CD spectrophotometer in a 0.1 cm path length quartz cuvette, using temperature/wavelength mode. Spectra were collected at 20 °C over a wavelength range from 200 to 250 nm. Apo- and holo-MPP1 were prepared at 4 µM and 3.8 µM, respectively, for 200-250 nm window and 0.5 mM for the visible region in 50 mM MES, 150 mM NaCl pH 7.5.

**Preparation of Anhydrous Ethereal Monochloramine.** $NH_4Cl$ (3 g, 56 mmol) in ether (110 mL) was cooled to −5 °C, and concentrated $NH_4OH$ (4.7 mL) was added via pipet. Commercial bleach (Clorox, 72 mL) was then added via addition funnel over 15 min. The mixture was stirred for 15 min, the layers were separated, and the organic layer was washed with brine (1 × 35 mL). The organic layer was dried over powdered $CaCl_2$ in a freezer for 1 h and stored at -80 °C. Approximate concentration is 0.15 M.[4]

**Synthesis of N-Chloro-2-phenyl aziridine**

Excess NaOCl (475 mM, 250 µL) was added to 2-phenylaziridine (23 mM) at room temperature in 1 mL MES buffer and mixed for 5 minutes. The colorless solution became a white turbid solution during this time. The solution was then extracted with EtOAc (1 mL x 3) and dried over $Na_2SO_4$. The solvent was evaporated and redissolved in EtOAc and stored at 4 ˚C. Yield = 87 % analyzed by GC-MS.

**Preparation of Resonance Raman Samples**

**a) Raman Sample Preparation for $Mn^V(^{14/15}N)$ DPP**

Crystalline Mn(V)(N)(DPP) (1-2 mg, MW = 529 g $mol^{-1}$) was dissolved in dichloromethane (2.0 mL) to give an approximately 1 mM solution. An aliquot (400 µL) of this solution was transferred to a clean, dry NMR tube, which was then sealed and purged with argon for 5 min. After purging, the sample was rapidly frozen by immersion in liquid nitrogen. To confirm the nature of the species in solution, an aliquot (1-2 µL) of the Mn(V)(N)(DPP) stock solution was transferred into a UV-vis cuvette containing dichloromethane (2.0 mL) and mixed thoroughly. The UV-vis absorption spectrum was recorded, and the absorbance at 407 nm was measured ($\varepsilon_{407} = 5.1 \times 10^4$ $M^{-1}$ $cm^{-1}$, path length = 1 cm). The concentration of Mn(V)(N)(DPP) in the diluted sample was calculated using the Beer-Lamberts law, and the concentration of the stock solution was back-calculated based on the dilution factor.

**b) Raman sample Preparation for Mn(V)($^{15}N$) MPP1**

$^{15}NH_4Cl$ (375 mg) was dissolved in 5 ml ice cold water containing NaOH (580 mg). This solution was mixed for 5 minutes at 0 ˚C and transferred to a falcon tube containing Mn(III)MPP1 (0.23 mM, 6 mL MES buffer). A 15 % sodium hypochlorite solution (3 ml) was added, and the reaction was stirred an additional 15 minutes, resulting in a purple solution. This sample was concentrated to 500 µL and was transferred to a clean, dry NMR tube, which was then sealed and purged with argon for 5 min. After purging, the sample was rapidly frozen by immersion in liquid nitrogen.

**Preparation of H64A MPP1**

The gene coding for the protein sequence of MPP1 H64A was ordered from Genscript, which was cloned into the pET-11a plasmid. The geneblock encoding H64A MPP1 is below:

ATGCATCACCACCACCACCACGAGAACCTGTACTTCCAAAGCAGCGAGAAAGAGG
AGCTGTTTGAAAAGCTGAAACAGACCGCGGATGAGGCGGTGCAGCTGTTCCAACG
TCTGCGTGAAATCTTTGACAAGGGTGACGATGACAGCTTCGAACAGGTTCTGGAG
GAACTGGAGGAAGCGCTGCAGAAAGCGCGTCAACTGGCGGATCAGGGTCGTAAG
AAAGGCCTGCTGACCAGCGAGGCGGCGAAGCAGGGTGATCAATTTGTTCAACTGT
TCCAACGTTTTCGTGAAGCGTGGGACAAGGGCGATAAAGACAGCCTGGAGCAAAT
CCTGGAGGAACTGGAACAGGTGGCGCAAAAAGCGGTTGAGCTGGGCCTGAAGAT
TCTGAAAACCCAGTAA

H64A MPP1 was expressed and purified as previously reported for MPP1.[6]

**Spectroscopic Characterizations**

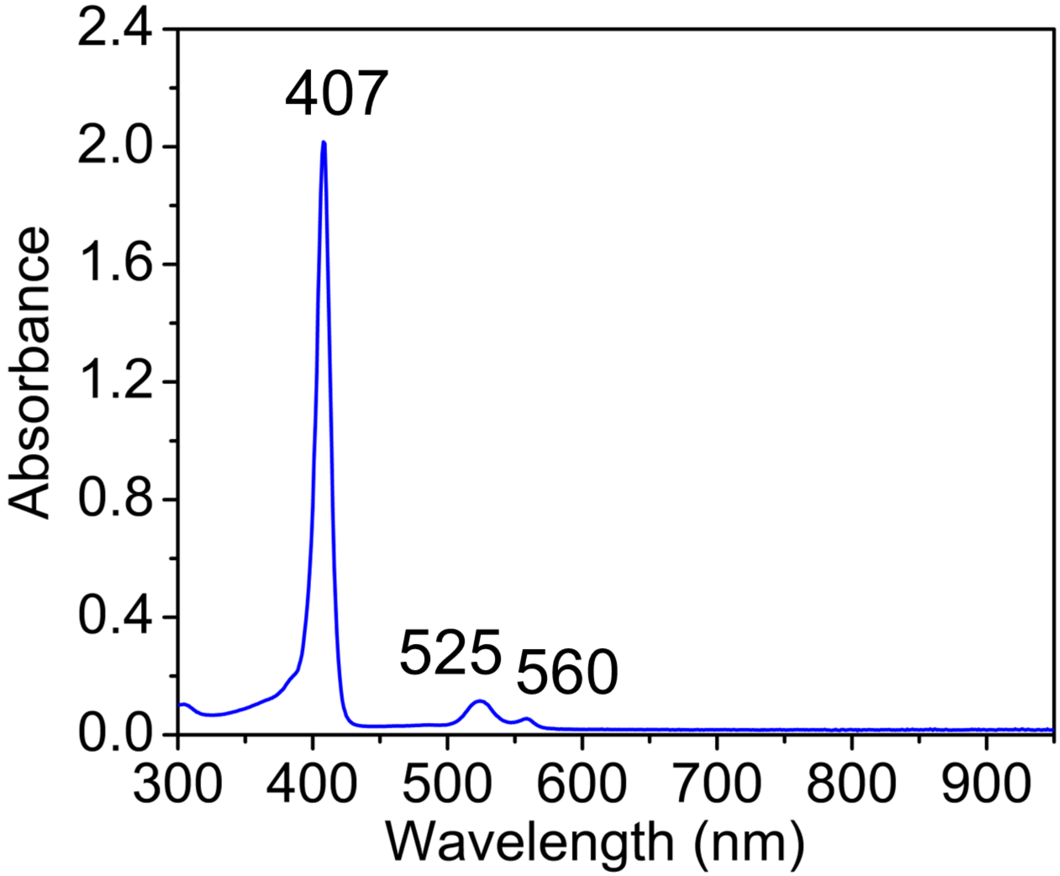


**Figure S1**. UV-vis of Mn(V)(N)-DPP (40 μM) cofactor in dichloromethane.

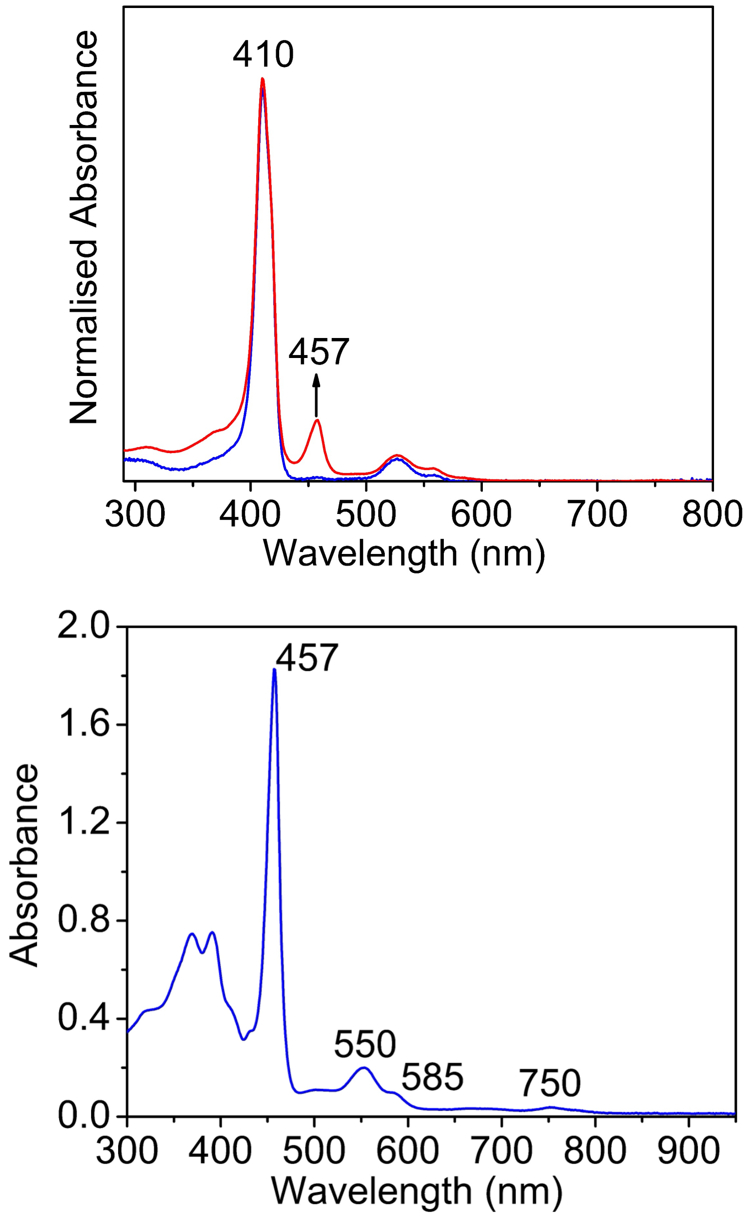


**Figure S2**. (a) UV-vis of crystalline Mn(V)(N)-DPP cofactor in DMSO (blue spectrum). An aliquot was taken from the concentrated Mn(N)-DPP solution (0.5 M in DMSO) showing slow decay after 2 days to $Mn^{III}$DPP as seen by the growth of peaks at 457 nm (red spectrum). (b) UV-vis of authentic $Mn^{III}$DPP(Cl) in DMSO (blue spectrum) showing Soret band at 457 nm matching with the decomposition product of Mn(V)(N) species.

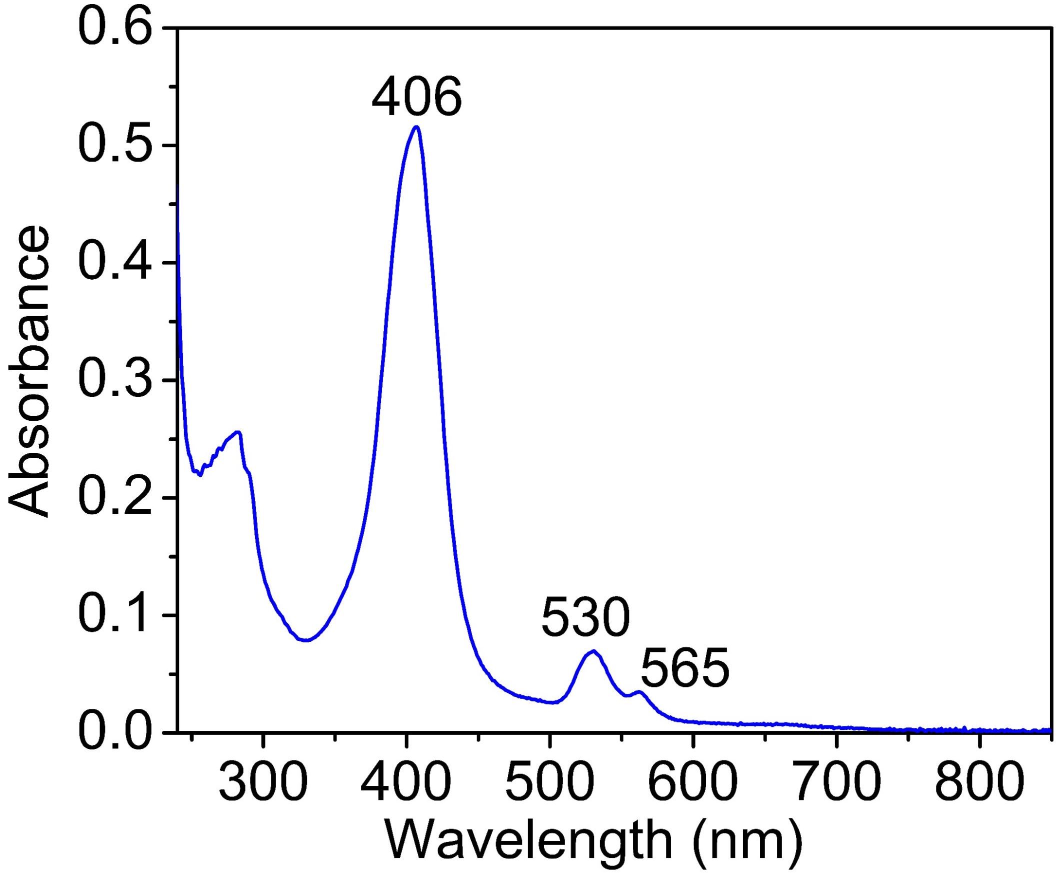


**Figure S3**. UV-vis of spectrum obtained after the addition of Mn(V)(N)-DPP to apo-MPP1 in MES buffer at 25 °C.

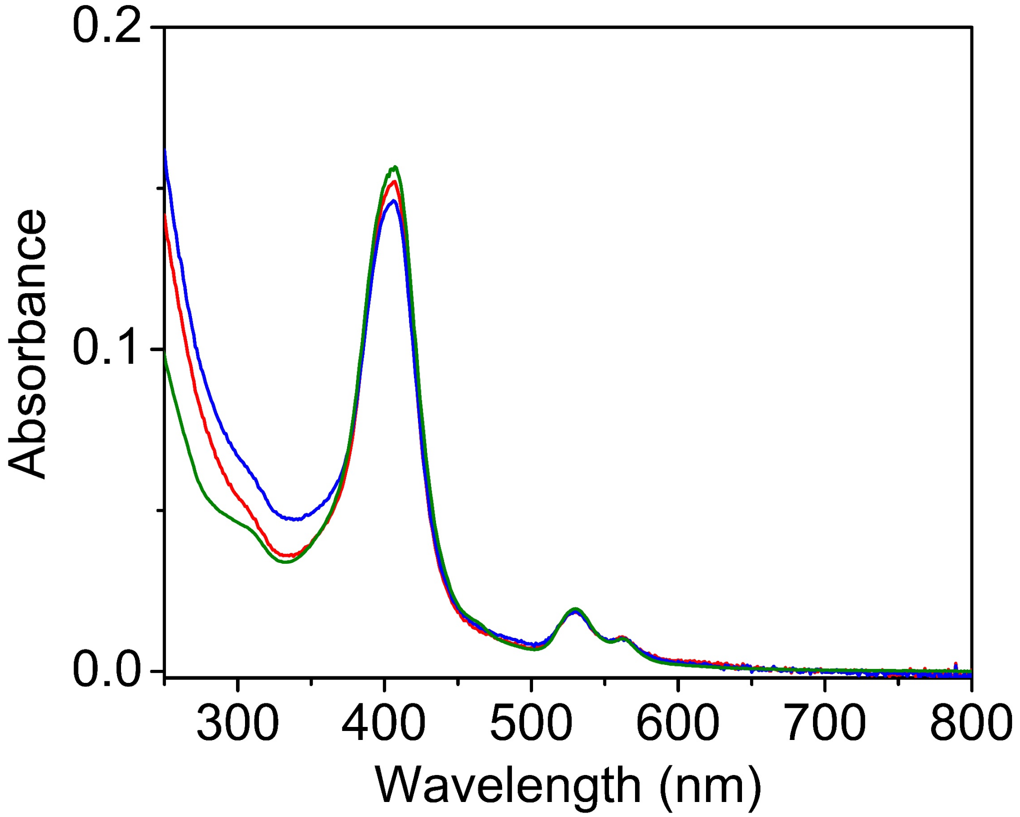


**Figure S4**. UV-vis spectra demonstrating the stability of Mn(V)(N)-MPP1 over time. The spectrum recorded on day 1 is shown in red, day 2 in blue, and after three months in green. The near-identical spectral features over this period indicate long-term stability of the Mn(V)-nitrido species inside the MPP1 protein.

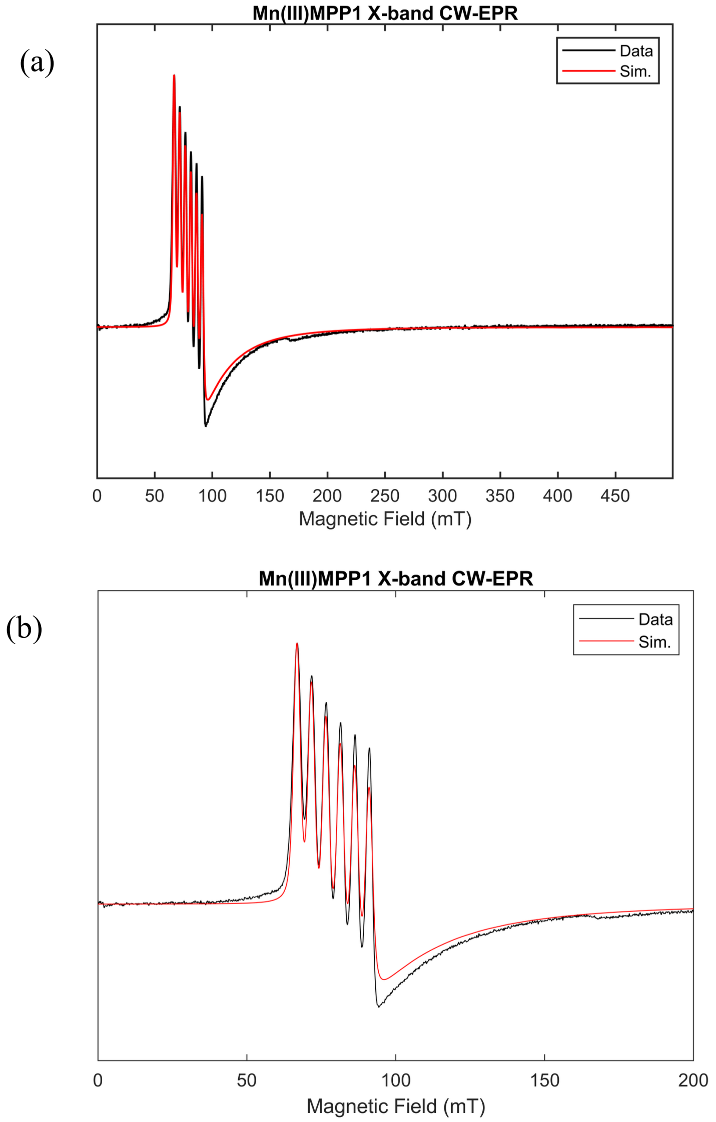


**Figure S5**. (a) X-band parallel-mode EPR spectrum of Mn(III)MPP1 recorded at 5 K. Mn(III)MPP1 is an integer-spin, non-Kramers ion and exhibits features consistent with an $S = 2$ ground state. (b) Well-resolved hyperfine splitting arising from coupling to the $^{55}$Mn nucleus ($I = 5/2$) is observed and a zoomed image is shown. The experimental spectrum (black) is well reproduced by simulation (red) using the Easyspin[1] Matlab toolbox with parameters of $g = 2.011$, zero-field splitting $D = \pm$ 2.43 cm$^{-}$ with rhombicity $E/D = 0.12$, and hyperfine coupling $A(^{55}\mathrm{Mn})$ = 136.3 MHz. The magnitude of the zero field splitting and hyperfine coupling is consistent with reported values for other Mn(III) complexes.[5] Acquisition Parameters: microwave power = 8.7 mW, field modulation amplitude = 0.8 mT, conversion time = 10.24 ms.

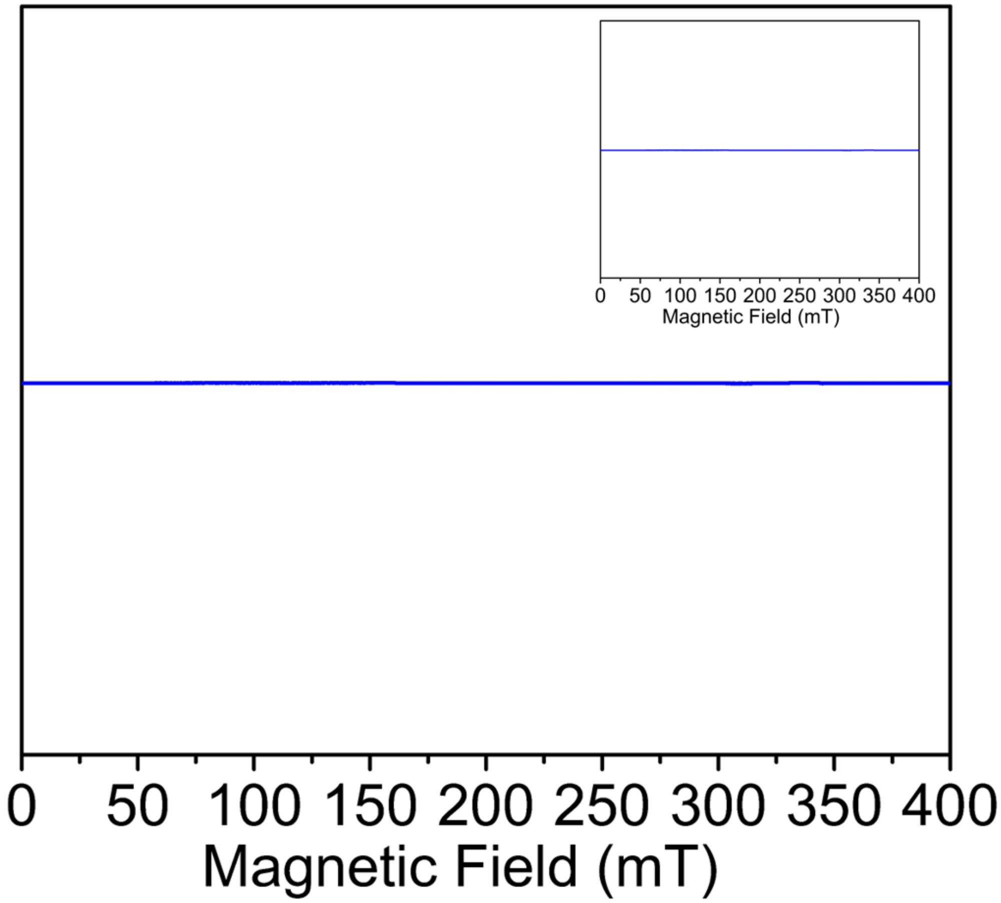


**Figure S6**. X-band EPR spectrum of the Mn(V)(N)-MPP1 species in parallel and perpendicular mode(inset) showing no detectable signal, consistent with a diamagnetic S = 0 ground state.

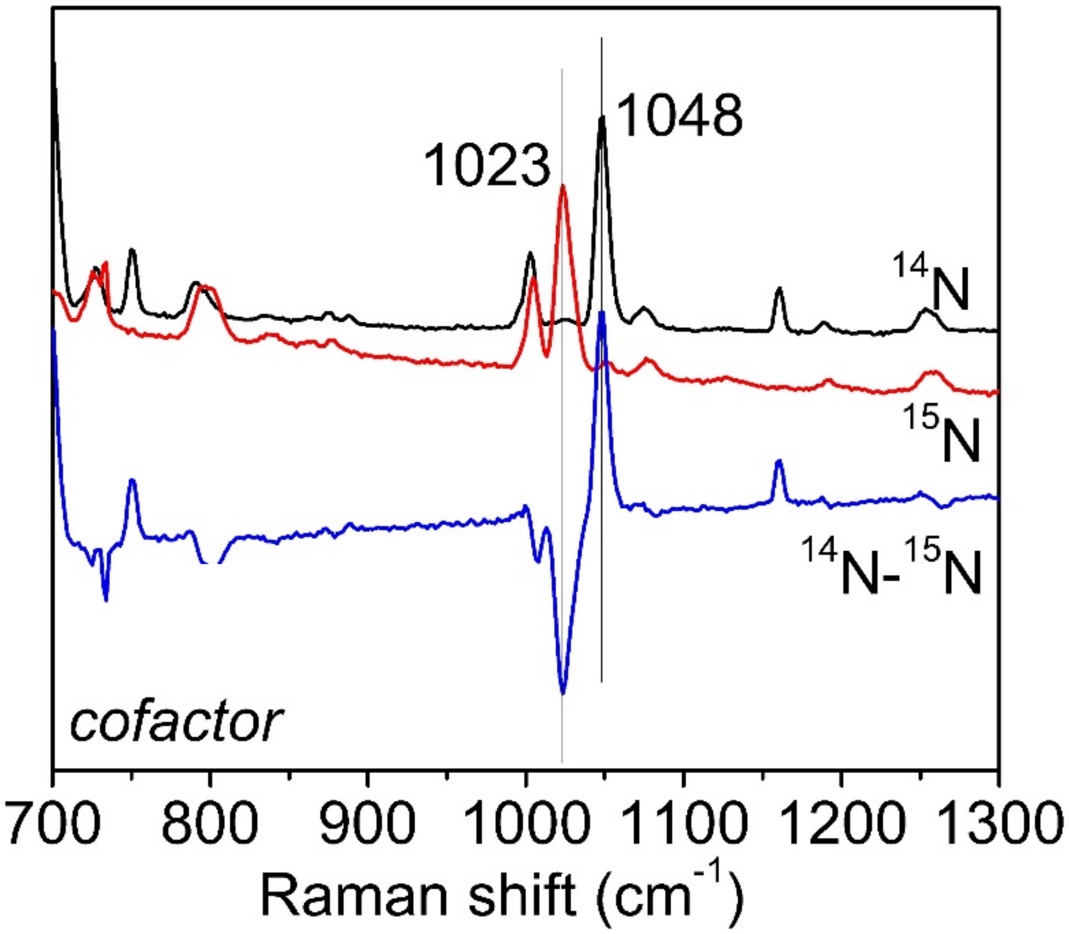


**Figure S7.** Low-temperature rR spectra of cofactor Mn($^{14}$N)(DPP) (black), Mn($^{15}$N)(DPP) (red) in DCM obtained with a 406.7-nm laser excitation (15 mW). The difference spectrum is shown in blue. An effective dielectric constant of ~9 for dichloromethane (DCM), the solvent used for RR experiments, provides a reasonable approximation of the similar electrostatic environment within hydrophobic protein interiors.[3]

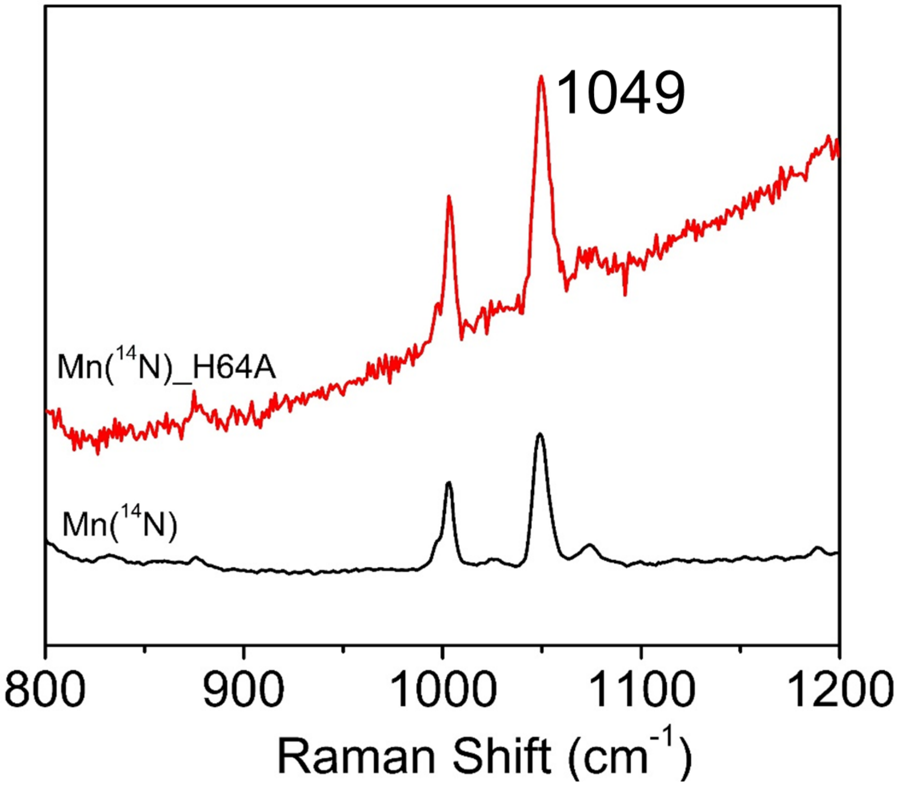


**Figure S8**. Low-temperature rR spectra of Mn(V)($^{14}$N)MPP1 (black), Mn(V)($^{14}$N)MPP1-H64A (red) in MES buffer obtained with a 406.7-nm laser excitation (8 mW) showing the peak at 1049 cm$^{-1}$ consistent with Mn-N stretching frequency.

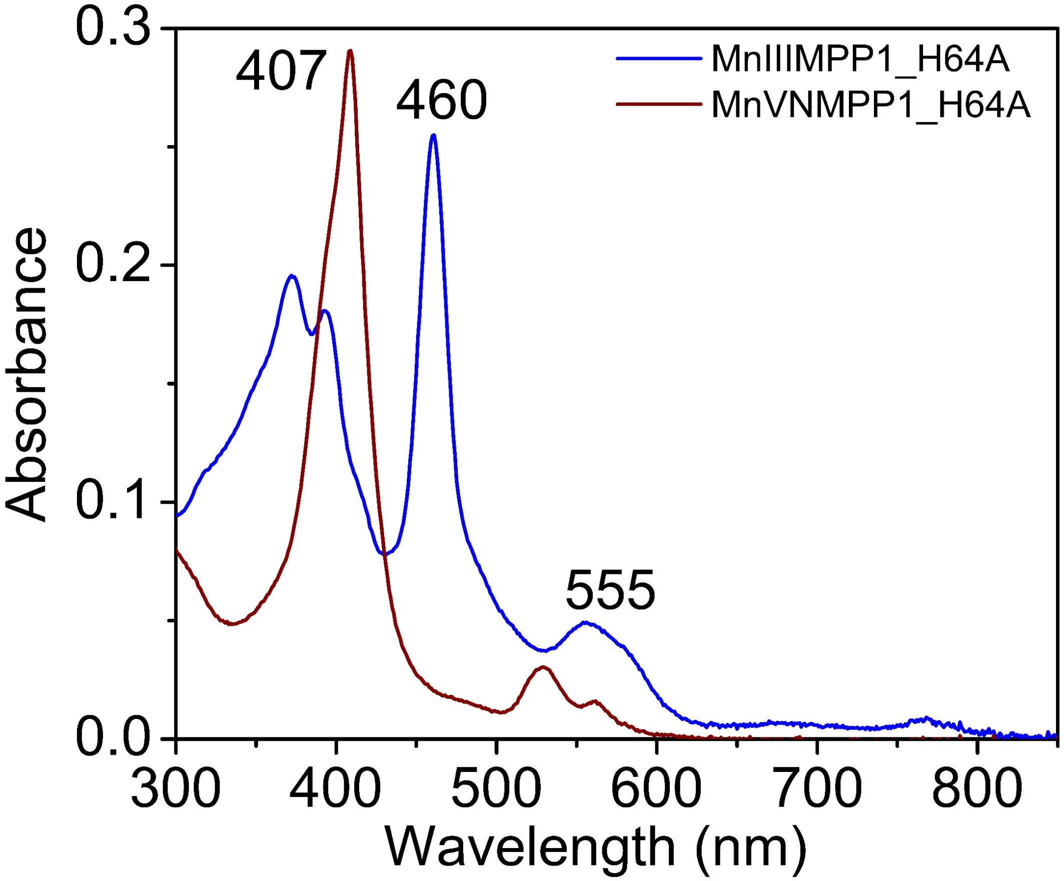


**Figure S9**. UV-vis spectra of Mn(III)MPP1 H64A and Mn(V)(N)MPP1 H64A highlighting the spectral changes from 460 to 407 nm associated with oxidation to the Mn(V)-nitrido species. Mutation of the axial histidine to alanine significantly alters the Q-band features of Mn(III)MPP1, consistent with strong axial coordination in the Mn(III) state. In contrast, no measurable changes are observed in the UV-vis features of Mn(V)(N)MPP1 upon the H64A mutation, indicating weak or negligible axial histidine coordination in the Mn(V)-nitrido state.

(a)

(b)

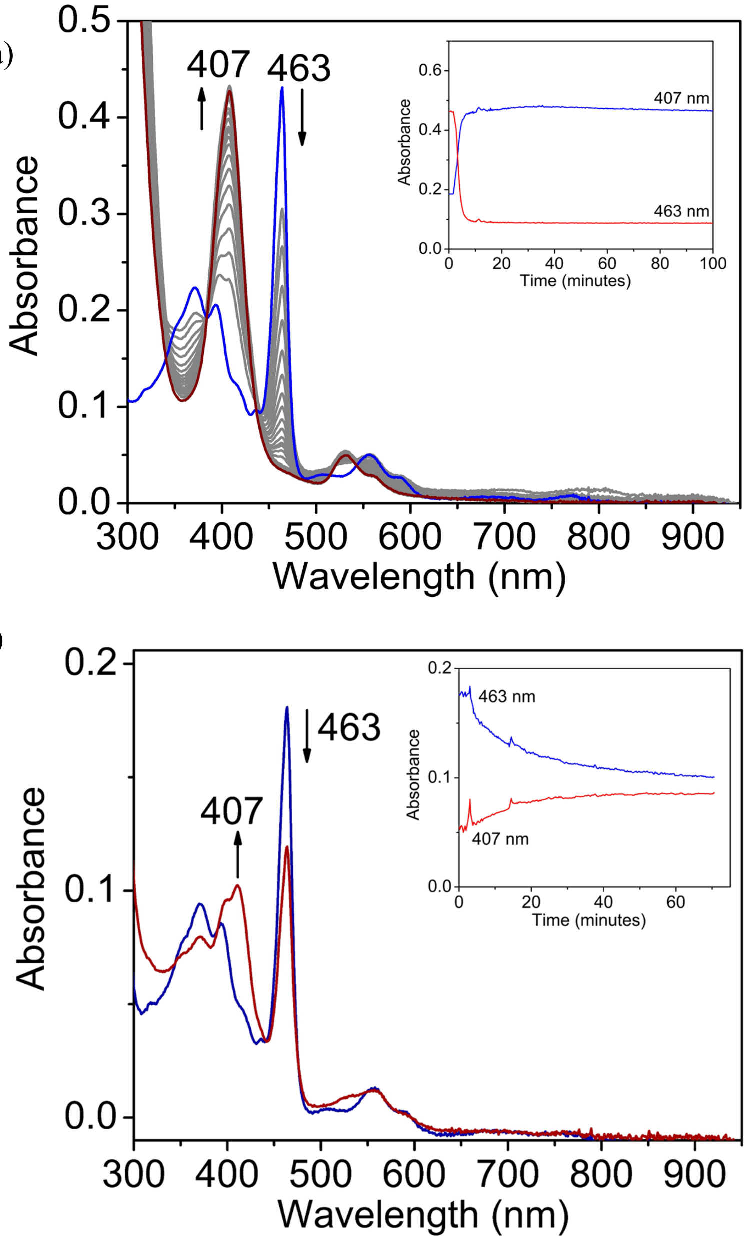


**Figure S10**. (a) UV-vis absorption spectra collected during the reaction of Mn(III)(MPP1) (blue spectrum) with $NH_2Cl$ (1.1 equivalent) to form Mn(V)(N)-MPP1(red spectrum) at RT within five minutes. The inset shows the absorption versus time curve at 407 and 463 nm during the reaction. Grey lines correspond to a time interval of 15 s. (b) UV-vis spectral changes observed during the reaction of Mn(III)MPP1 with $NH_2Cl$ (1.1 equivalent) at room temperature in the presence of excess substrate, showing the gradual formation of Mn(V)N-MPP1 over time. Under these conditions, Mn(V)N-MPP1 accumulates slowly and reaches ~50% yield. The diminished accumulation relative to substrate-free reactions suggests that a precursor intermediate to Mn(V)N-MPP1 undergoes competitive reaction with the substrate, thereby suppressing full nitrido formation.

**Nitrogen Atom Transfer Experiment of Mn(V)N-MPP1**

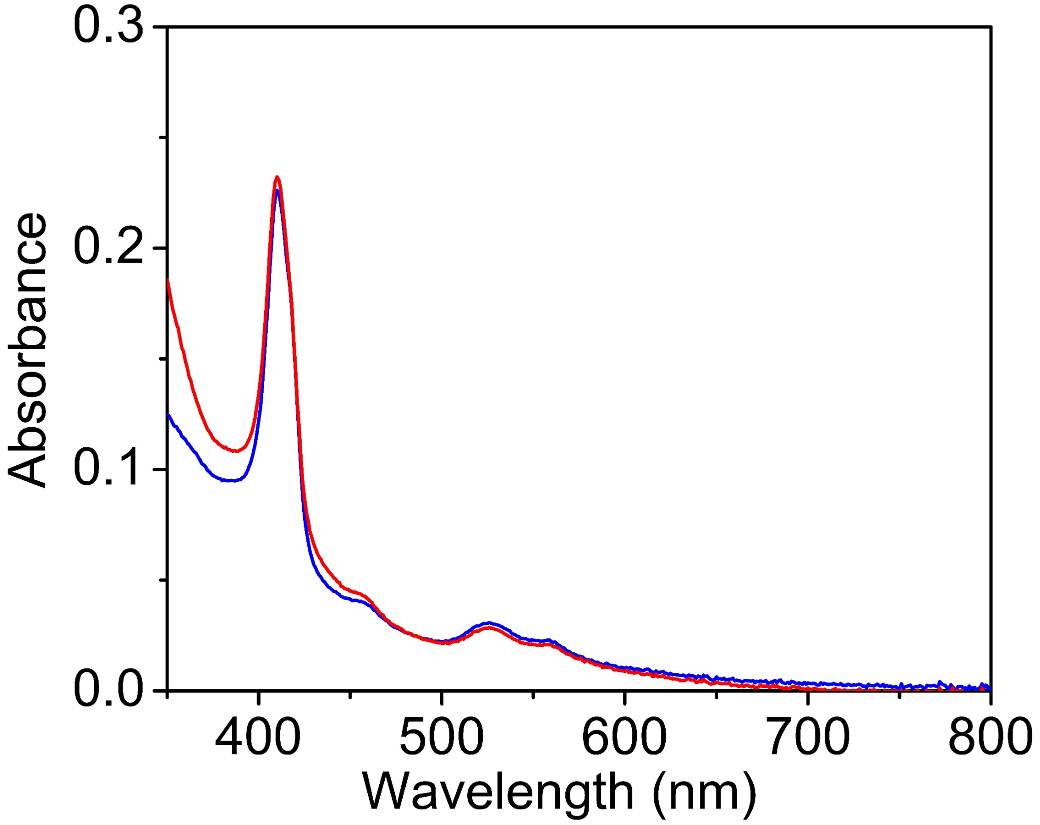


**Figure S11**. Reaction of Mn(V)(N)-MPP1(initial blue spectrum) with styrene at RT showing no reaction (red spectrum).

**Reactivity Studies - Mn(III)MPP1 during catalytic conditions**

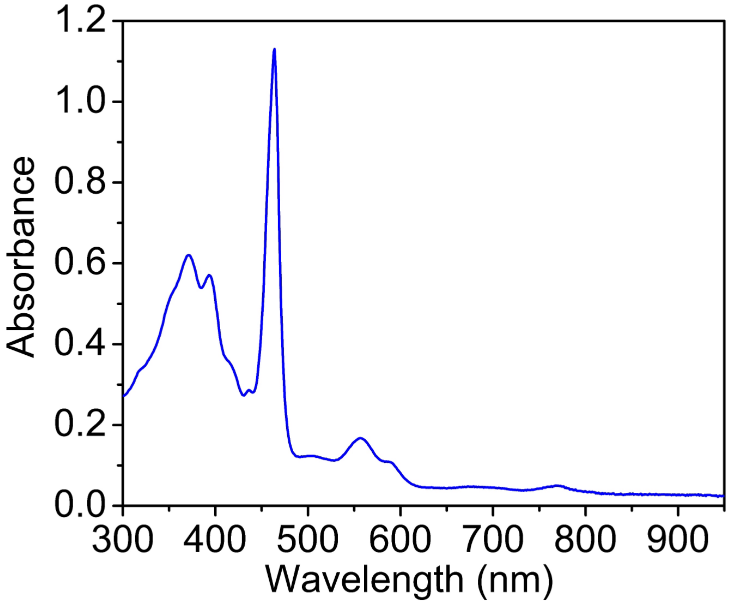


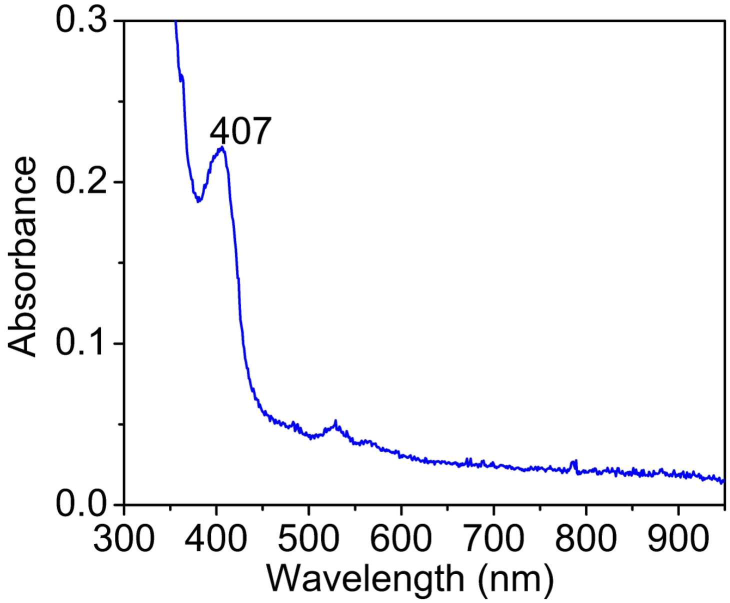


**Figure S12**. UV-vis spectra of (top) Mn(III)-MPP1 (16 μM) in the presence of styrene in MES buffer at room temperature and (bottom) the same sample recorded 5 min after addition of $NH_4OH$ and NaOCl, showing minimal formation of the Mn(V)(N)-MPP1 species during catalytic cycle.

**Mn(V)NMPP1 with styrene – GC-MS**

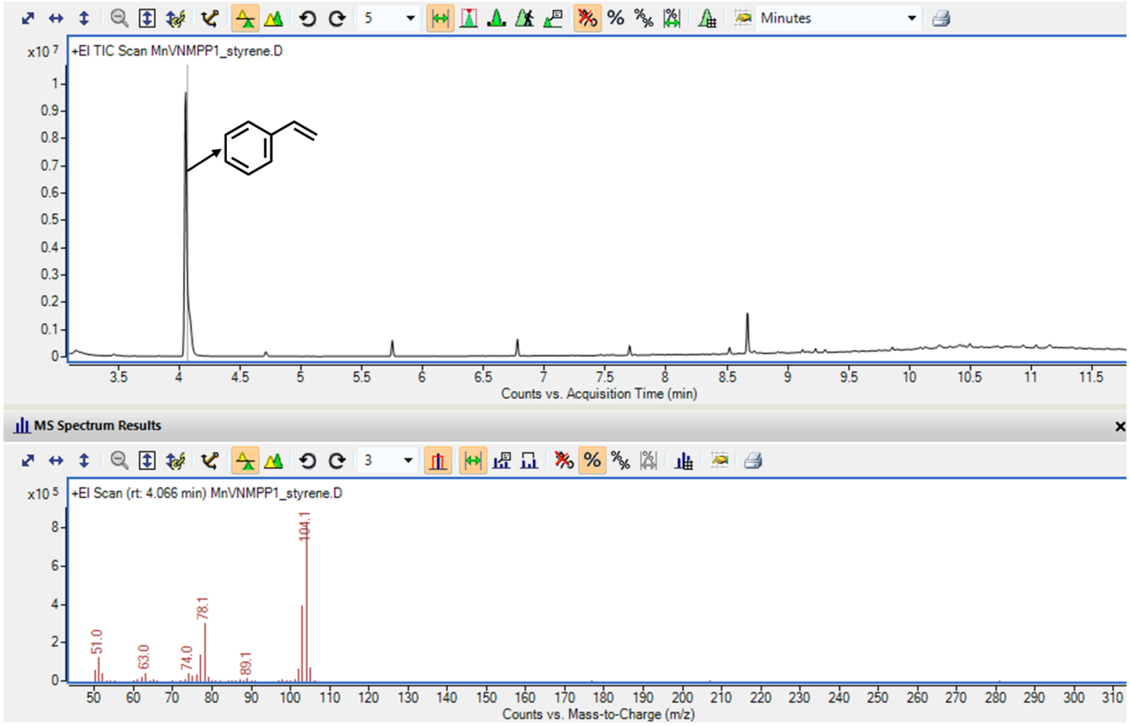


**Figure S13**. GC-MS spectrum for the reaction of Mn(V)(N)MPP1 with styrene at room temperature showing no formation of 2-phenylaziridine or N-Chloro-2-phenylaziridine products at 5.98 and 6.3 minutes, respectively. The styrene peak at a retention time of 4.1 minutes can be clearly observed.

**Table S1. Bond Parameters of Mn(V)N-DPP**

| Atom1 | Atom2 | Length |
|---|---|---|
| N2 | C1 | 1.385(3) |
| N2 | Mn1 | 1.972(2) |
| N2 | C14 | 1.378(3) |
| N2 | Mn1 | 2.088(2) |
| N3 | C9 | 1.386(3) |
| N3 | C12 | 1.380(3) |
| N3 | Mn1 | 2.058(2) |
| N3 | Mn1 | 1.983(2) |
| C1 | C2 | 1.396(4) |
| C1 | C16 | 1.435(4) |
| C2 | C3 | 1.497(4) |
| C2 | C9 | 1.391(4) |
| C3 | C4 | 1.392(4) |
| C3 | C8 | 1.395(4) |
| C4 | C5 | 1.390(4) |
| C5 | C6 | 1.386(4) |
| C6 | C7 | 1.387(4) |
| C7 | C8 | 1.389(4) |
| N3 | C12 | 1.380(3) |
| N3 | Mn1 | 2.058(2) |
| C1 | C2 | 1.396(4) |
| C2 | C3 | 1.497(4) |
| C2 | C9 | 1.391(4) |
| C3 | C4 | 1.392(4) |
| C3 | C8 | 1.395(4) |
| C4 | C5 | 1.390(4) |
| C5 | C6 | 1.386(4) |
| C6 | C7 | 1.387(4) |
| C7 | C8 | 1.389(4) |
| C9 | C10 | 1.437(4) |
| C10 | C11 | 1.352(4) |
| C11 | C12 | 1.434(4) |
| C12 | C13 | 1.379(4) |
| C13 | C14 | 1.382(4) |
| C14 | C15 | 1.430(4) |
| C15 | C16 | 1.356(4) |
| Mn1 | N1 | 1.534(4) |
| C9 | C10 | 1.437(4) |
| C10 | C11 | 1.352(4) |
| C11 | C12 | 1.434(4) |
| C12 | C13 | 1.379(4) |
| C13 | C14 | 1.382(4) |
| C14 | C15 | 1.430(4) |
| C14 | N2 | 1.378(3) |
| C15 | C16 | 1.356(4) |
| C16 | C1 | 1.435(4) |
| Mn1 | N1 | 1.534(4) |
| Mn1 | N2 | 2.088(2) |
| Mn1 | N3 | 1.983(2) |
| Mn1 | Mn1 | 0.778(1) |
| Mn1 | N1 | 2.305(4) |
| N1 | Mn1 | 2.305(4) |
| N2 | C1 | 1.385(3) |
| N2 | Mn1 | 1.972(2) |
| N3 | C9 | 1.386(3) |

**Table S2. Bond Angles of Mn(V)(N)-DPP**

| Atom1 | Atom2 | Atom3 | Angle |
|---|---|---|---|
| C1 | N2 | Mn1 | 128.7(2) |
| C1 | N2 | C14 | 104.8(2) |
| C1 | N2 | Mn1 | 126.9(2) |
| Mn1 | N2 | C14 | 124.4(2) |
| Mn1 | N2 | Mn1 | 21.84(4) |
| C14 | N2 | Mn1 | 127.2(2) |
| C9 | N3 | C12 | 105.1(2) |
| C9 | N3 | Mn1 | 125.6(2) |
| C9 | N3 | Mn1 | 129.6(2) |
| C12 | N3 | Mn1 | 128.0(2) |
| C12 | N3 | Mn1 | 123.3(2) |
| Mn1 | N3 | Mn1 | 22.09(4) |
| N2 | C1 | C2 | 125.2(2) |
| N2 | C1 | C16 | 110.2(2) |
| C2 | C1 | C16 | 124.5(3) |
| C1 | C2 | C3 | 119.2(2) |
| C1 | C2 | C9 | 123.0(3) |
| C3 | C2 | C9 | 117.9(2) |
| C2 | C3 | C4 | 121.3(2) |
| C2 | C3 | C8 | 119.9(2) |
| C4 | C3 | C8 | 118.8(3) |
| C3 | C4 | H4 | 119.8 |
| C3 | C4 | C5 | 120.5(3) |
| H4 | C4 | C5 | 119.8 |
| C4 | C5 | H5 | 119.9 |
| C4 | C5 | C6 | 120.3(3) |
| H5 | C5 | C6 | 119.9 |
| C5 | C6 | H6 | 120.1 |
| C5 | C6 | C7 | 119.8(3) |
| H6 | C6 | C7 | 120.1 |
| C6 | C7 | H7 | 120.1 |
| C6 | C7 | C8 | 119.8(3) |
| H7 | C7 | C8 | 120.1 |
| C3 | C8 | C7 | 120.8(3) |
| C3 | C8 | H8 | 119.6 |
| C7 | C8 | H8 | 119.6 |
| N3 | C9 | C2 | 125.8(2) |
| N3 | C9 | C10 | 110.1(2) |
| C2 | C9 | C10 | 124.1(3) |
| C9 | C10 | H10 | 126.3 |
| C9 | C10 | C11 | 107.3(2) |
| H10 | C10 | C11 | 126.4 |
| C10 | C11 | H11 | 126.5 |
| C10 | C11 | C12 | 106.9(2) |
| H11 | C11 | C12 | 126.5 |
| N3 | C12 | C11 | 110.6(2) |
| N3 | C12 | C13 | 125.3(2) |
| C11 | C12 | C13 | 124.1(3) |
| C12 | C13 | H13 | 117.2 |
| C12 | C13 | C14 | 125.7(3) |
| H13 | C13 | C14 | 117.2 |
| C13 | C14 | C15 | 124.0(3) |
| C13 | C14 | N2 | 124.9(2) |
| C15 | C14 | N2 | 111.2(2) |
| C14 | C15 | H15 | 126.7 |
| C14 | C15 | C16 | 106.5(2) |
| H15 | C15 | C16 | 126.7 |
| C15 | C16 | H16 | 126.3 |
| C15 | C16 | C1 | 107.3(2) |
| H16 | C16 | C1 | 126.4 |
| N2 | Mn1 | N3 | 87.46(9) |
| N2 | Mn1 | N1 | 100.0(2) |
| N2 | Mn1 | N2 | 158.2(1) |
| N2 | Mn1 | N3 | 91.64(9) |
| N2 | Mn1 | Mn1 | 87.5(1) |
| N2 | Mn1 | N1 | 82.5(1) |
| N3 | Mn1 | N1 | 101.5(2) |
| N3 | Mn1 | N2 | 86.34(9) |
| N3 | Mn1 | N3 | 157.9(1) |
| N3 | Mn1 | Mn1 | 73.5(1) |
| N3 | Mn1 | N1 | 76.8(1) |
| N1 | Mn1 | N2 | 101.7(2) |
| N1 | Mn1 | N3 | 100.4(2) |
| N1 | Mn1 | Mn1 | 170.9(2) |
| N1 | Mn1 | N1 | 176.9(2) |
| N2 | Mn1 | N3 | 86.36(9) |
| N2 | Mn1 | Mn1 | 70.7(1) |

| | | | |
|---|---|---|---|
| N2 | Mn1 | N1 | 75.7(1) |
| N3 | Mn1 | Mn1 | 84.4(1) |
| N3 | Mn1 | N1 | 81.2(1) |
| Mn1 | Mn1 | N1 | 6.0(1) |
| Mn1 | N1 | Mn1 | 3.06(7) |
| C14 | N2 | Mn1 | 127.2(2) |
| C14 | N2 | C1 | 104.8(2) |
| C14 | N2 | Mn1 | 124.4(2) |
| Mn1 | N2 | C1 | 126.9(2) |
| Mn1 | N2 | Mn1 | 21.84(4) |
| C1 | N2 | Mn1 | 128.7(2) |
| Mn1 | N3 | C9 | 129.6(2) |
| Mn1 | N3 | C12 | 123.3(2) |
| Mn1 | N3 | Mn1 | 22.09(4) |
| C9 | N3 | C12 | 105.1(2) |
| C9 | N3 | Mn1 | 125.6(2) |
| C12 | N3 | Mn1 | 128.0(2) |
| C16 | C1 | N2 | 110.2(2) |
| C16 | C1 | C2 | 124.5(3) |
| N2 | C1 | C2 | 125.2(2) |
| C1 | C2 | C3 | 119.2(2) |
| C1 | C2 | C9 | 123.0(3) |
| C3 | C2 | C9 | 117.9(2) |
| C2 | C3 | C4 | 121.3(2) |
| C2 | C3 | C8 | 119.9(2) |
| C4 | C3 | C8 | 118.8(3) |
| C3 | C4 | H4 | 119.8 |
| C3 | C4 | C5 | 120.5(3) |
| H4 | C4 | C5 | 119.8 |
| C4 | C5 | H5 | 119.9 |
| C4 | C5 | C6 | 120.3(3) |
| H5 | C5 | C6 | 119.9 |
| C5 | C6 | H6 | 120.1 |
| C5 | C6 | C7 | 119.8(3) |
| H6 | C6 | C7 | 120.1 |
| C6 | C7 | H7 | 120.1 |
| C6 | C7 | C8 | 119.8(3) |
| H7 | C7 | C8 | 120.1 |
| C3 | C8 | C7 | 120.8(3) |
| C3 | C8 | H8 | 119.6 |
| C7 | C8 | H8 | 119.6 |

| | | | |
|---|---|---|---|
| N3 | C9 | C2 | 125.8(2) |
| N3 | C9 | C10 | 110.1(2) |
| C2 | C9 | C10 | 124.1(3) |
| C9 | C10 | H10 | 126.3 |
| C9 | C10 | C11 | 107.3(2) |
| H10 | C10 | C11 | 126.4 |
| C10 | C11 | H11 | 126.5 |
| C10 | C11 | C12 | 106.9(2) |
| H11 | C11 | C12 | 126.5 |
| N3 | C12 | C11 | 110.6(2) |
| N3 | C12 | C13 | 125.3(2) |
| C11 | C12 | C13 | 124.1(3) |
| C12 | C13 | H13 | 117.2 |
| C12 | C13 | C14 | 125.7(3) |
| H13 | C13 | C14 | 117.2 |
| N2 | C14 | C13 | 124.9(2) |
| N2 | C14 | C15 | 111.2(2) |
| C13 | C14 | C15 | 124.0(3) |
| C14 | C15 | H15 | 126.7 |
| C14 | C15 | C16 | 106.5(2) |
| H15 | C15 | C16 | 126.7 |
| C1 | C16 | C15 | 107.3(2) |
| C1 | C16 | H16 | 126.4 |
| C15 | C16 | H16 | 126.3 |
| N2 | Mn1 | N3 | 86.36(9) |
| N2 | Mn1 | Mn1 | 70.7(1) |
| N2 | Mn1 | N1 | 75.7(1) |
| N2 | Mn1 | N2 | 158.2(1) |
| N2 | Mn1 | N3 | 86.34(9) |
| N2 | Mn1 | N1 | 101.7(2) |
| N3 | Mn1 | Mn1 | 84.4(1) |
| N3 | Mn1 | N1 | 81.2(1) |
| N3 | Mn1 | N2 | 91.64(9) |
| N3 | Mn1 | N3 | 157.9(1) |
| N3 | Mn1 | N1 | 100.4(2) |
| Mn1 | Mn1 | N1 | 6.0(1) |
| Mn1 | Mn1 | N2 | 87.5(1) |
| Mn1 | Mn1 | N3 | 73.5(1) |
| Mn1 | Mn1 | N1 | 170.9(2) |
| N1 | Mn1 | N2 | 82.5(1) |
| N1 | Mn1 | N3 | 76.8(1) |

| | | | |
|---|---|---|---|
| N1 | Mn1 | N1 | 176.9(2) |
| N2 | Mn1 | N3 | 87.46(9) |
| N2 | Mn1 | N1 | 100.0(2) |
| N3 | Mn1 | N1 | 101.5(2) |
| Mn1 | N1 | Mn1 | 3.06(7) |

**Table S3. Reactivity studies summarizing preliminary catalytic activity and product yield determination.**

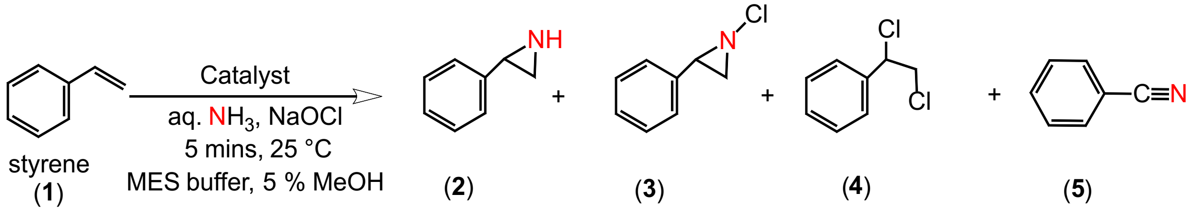


| Reaction conditions | 2 | 3 | 4 | 5 |
|---|---|---|---|---|
| Mn(III)MPP1 (25 uM) | not observed | 50 % | 30 % | n.d. |
| Mn(III)MPP1H64A (25 uM) | not observed | 15 % | 12 % | n.d. |
| Mn(V)NMPP1 | not observed | not observed | not observed | n.d. |
| No holo | not observed | 15 % | 1 % | n.d. |
| Mn(III)DPPCl cofactor | not observed | 15 % | 2 % | n.d. |

**Standard conditions**: Catalyst concentration = 25 µM, $NH_4OH$ = 740 mM, NaOCl = 475 mM, styrene = 10 mM in MeOH, 1 mL total volume, reaction time = 5 minutes. n.d. means not determined due to negligible amount.

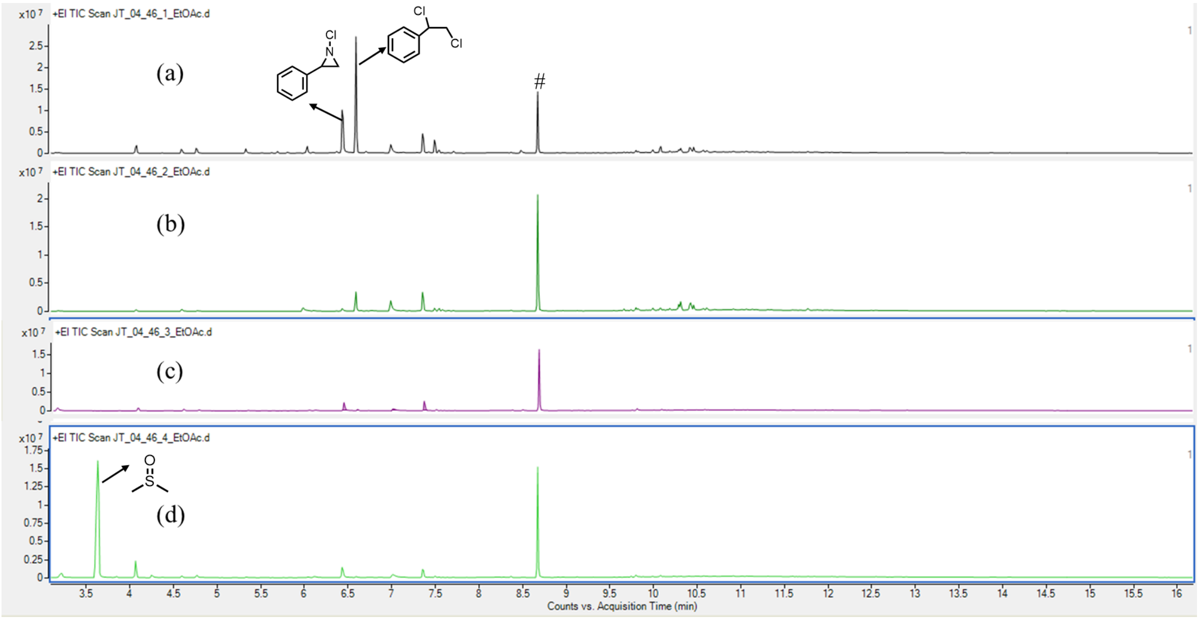


**Figure S14**. GC-MS traces observed under standard reaction conditions with (a) Mn(III)MPP1 (b) Mn(III)MPP1-H64A (c) No protein (d) Mn(III)DPP(Cl) cofactor-only. The (#) mark indicates unidentified impurity in the GC-MS column.

**Enantioselectivity Determination and UPC Analysis of N-Chloro Phenyl Aziridine Product.**

Conditions: 5% IPA/Hexanes, 1.0 mL/min, Injection volume - 5 μL

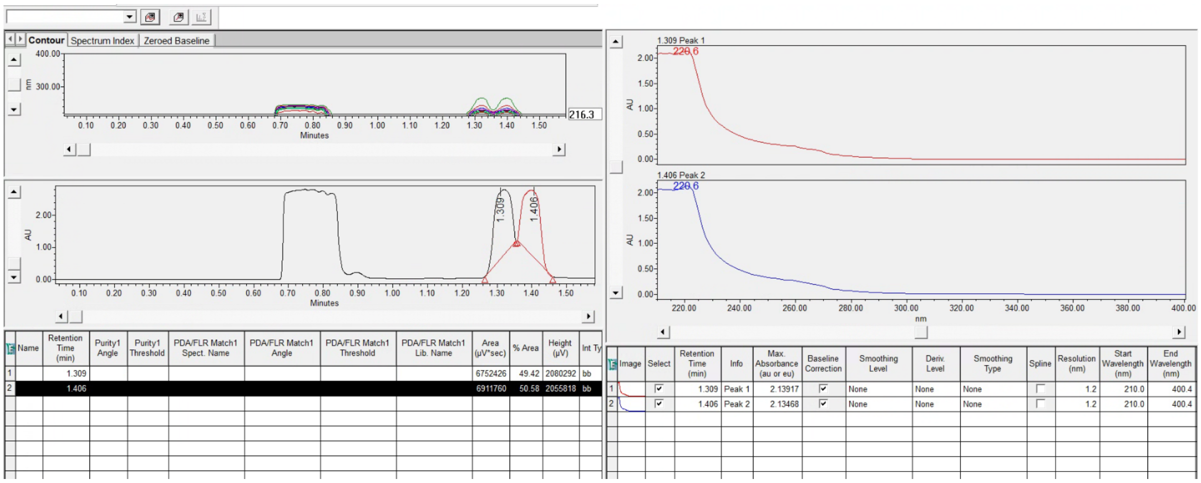


**Figure S15**. UltraPerformance Convergence Chromatography *(UPC²)* analysis of N-chloro-2-phenyl aziridine synthesized via the method described above, showing formation of a racemic product with two enantiomeric peaks observed at retention times of 1.31 and 1.41 min. The peak at 0.8 min corresponds to solvent EtOAc.

**Synthesis of N-Chloro phenyl aziridine using NaOCl**

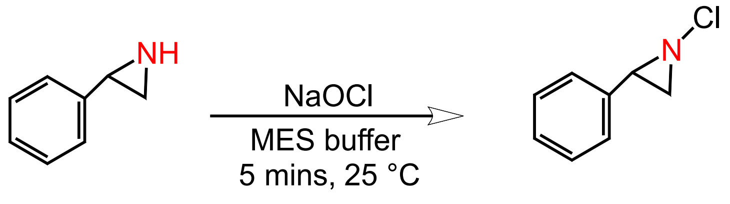


(**2**) (**3**)

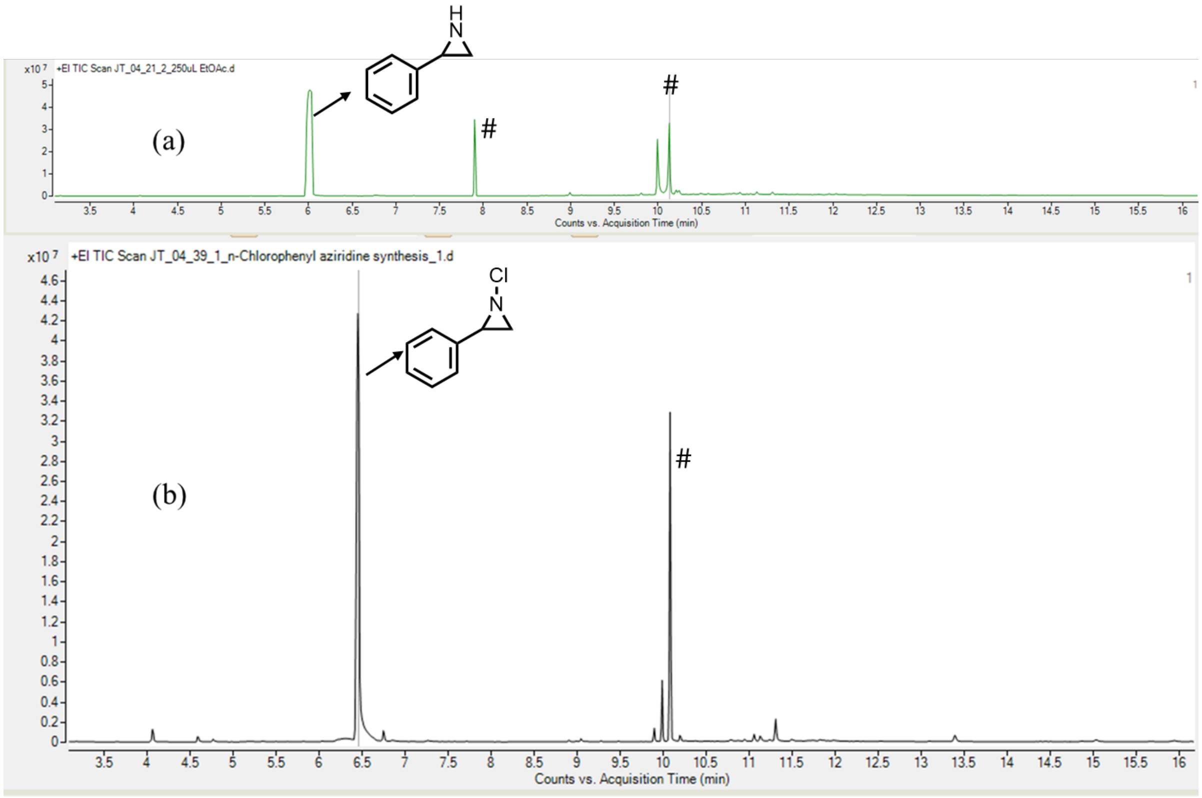


**Figure S16**. (a) GC-MS spectrum of commercially available 2-phenylaziridine (**2**) with retention time of 5.98 min (b) GC-MS spectrum of the reaction of 2-phenylaziridine with NaOCl in MES buffer, followed by extraction into ethyl acetate. Complete consumption of 2-phenyl aziridine (**2**) is observed. A new product peak appears at a retention time of 6.3 min and is assigned to N-chloro-2-phenyl aziridine, based on the characteristic 3:1 chlorine isotope pattern observed at $m/z$ = 153. The (#) corresponds to unidentified impurity peaks in the commercially available 2-Phenyl aziridine.

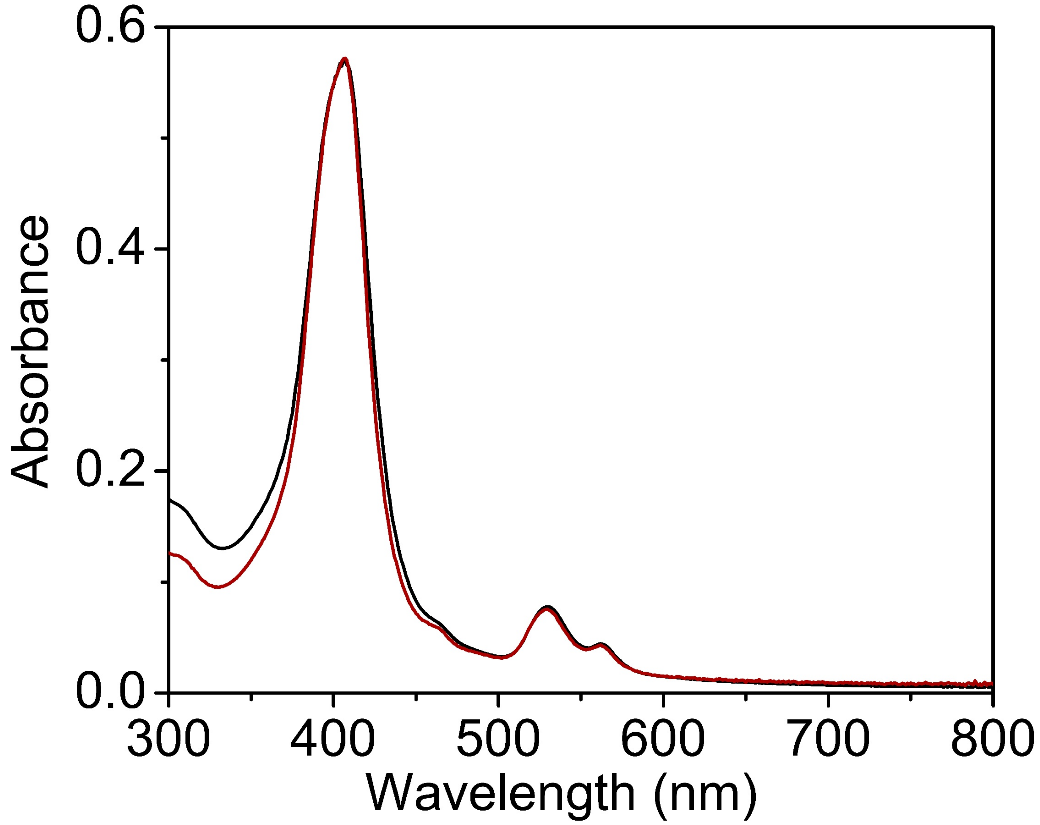


**Figure S17**. Reaction of Mn(V)(N)MPP1(initial black spectrum) with $Na_2S_2O_4$ (200 equivalent) at RT showing no reaction of the holo protein (final red spectrum).

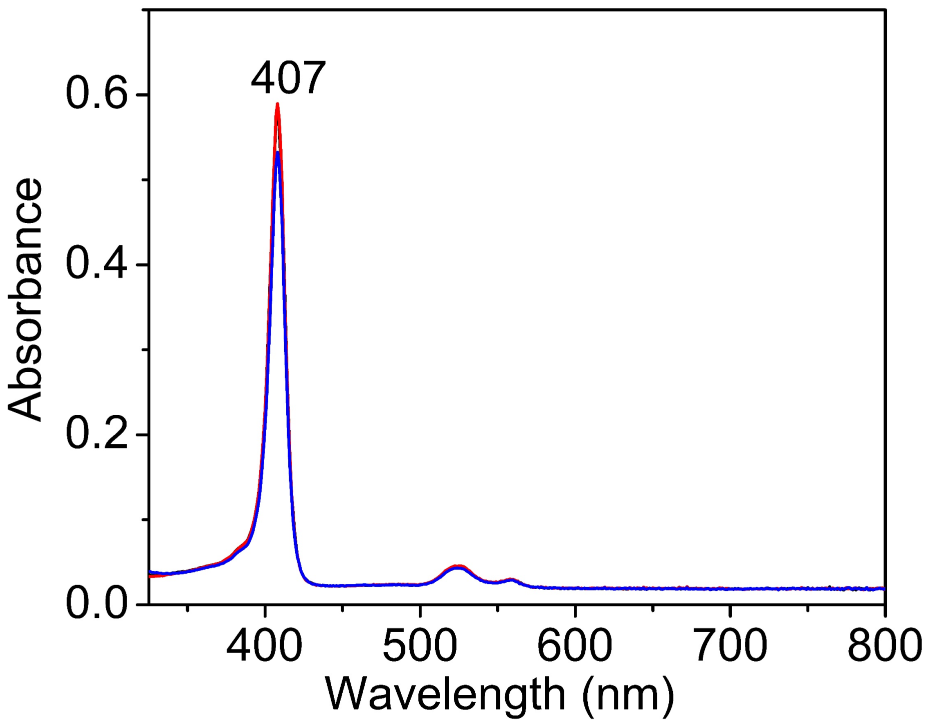


**Figure S18**. UV-vis changes upon addition of 1-methyl imidazole (1-MeIm, 2000 equivalent) Mn(V)(N)(DPP) cofactor in DCM. Red spectrum corresponds to initial Mn(V)(N)(DPP) cofactor, blue spectrum corresponds to after the addition of 1-MeIm axial ligand showing no change..

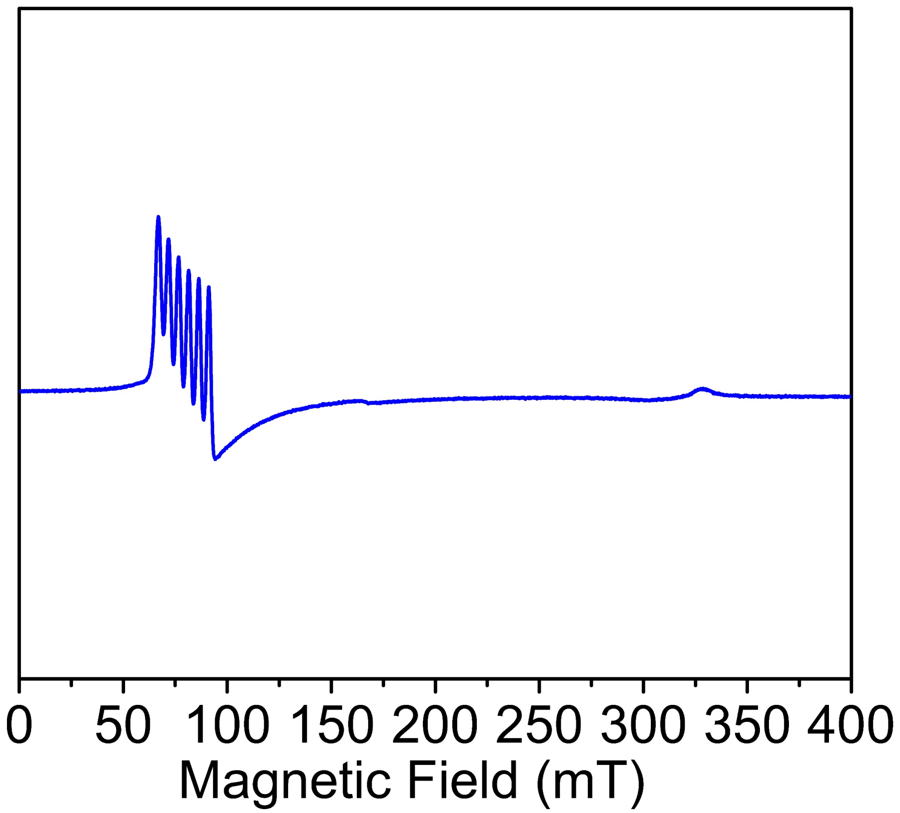


**Figure S19**. X-band parallel-mode EPR spectrum of Mn(III)-MPP1 in the presence of substrate styrene (10 mM) recorded at 5 K. The spectral features matched with the Mn(III)-MPP1 shown in **Figure S5**.

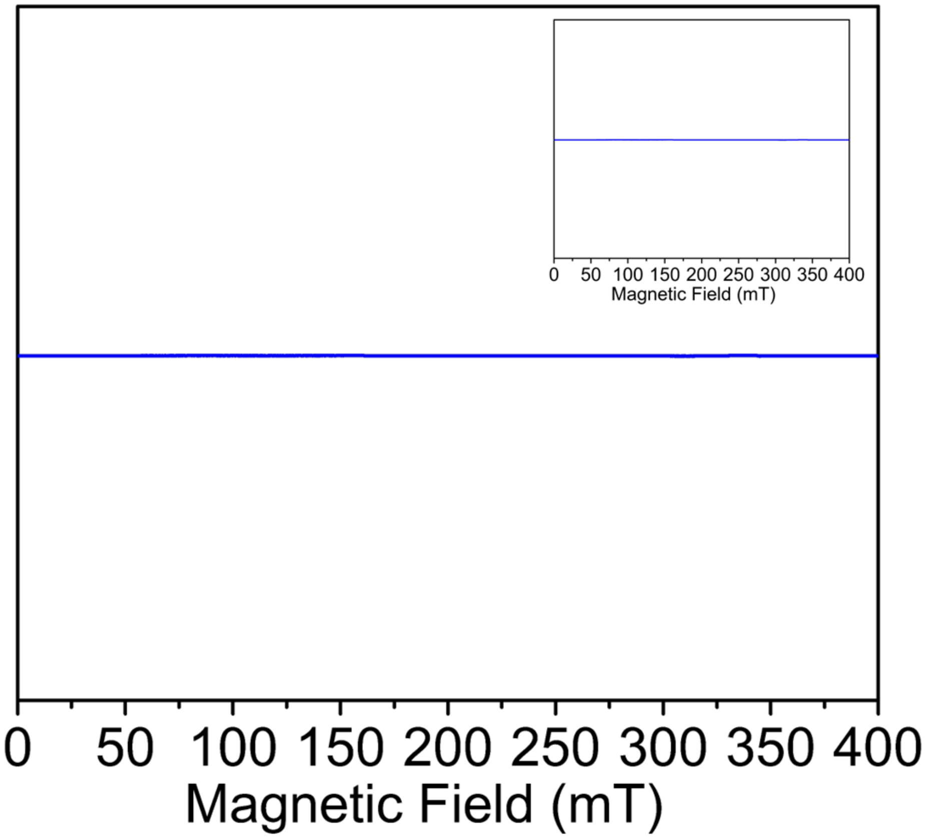


**Figure S20**. X-band parallel and perpendicular (inset) mode EPR spectrum of the reaction mixture obtained from Mn(III)-MPP1 treated with styrene, $NH_4OH$, and NaOCl, mixed for 5 min prior to freezing. The absence of an EPR signal indicates formation of an EPR-silent species, consistent with generation of the diamagnetic Mn(V)(N)-MPP1 intermediate on this timescale under catalytic conditions.

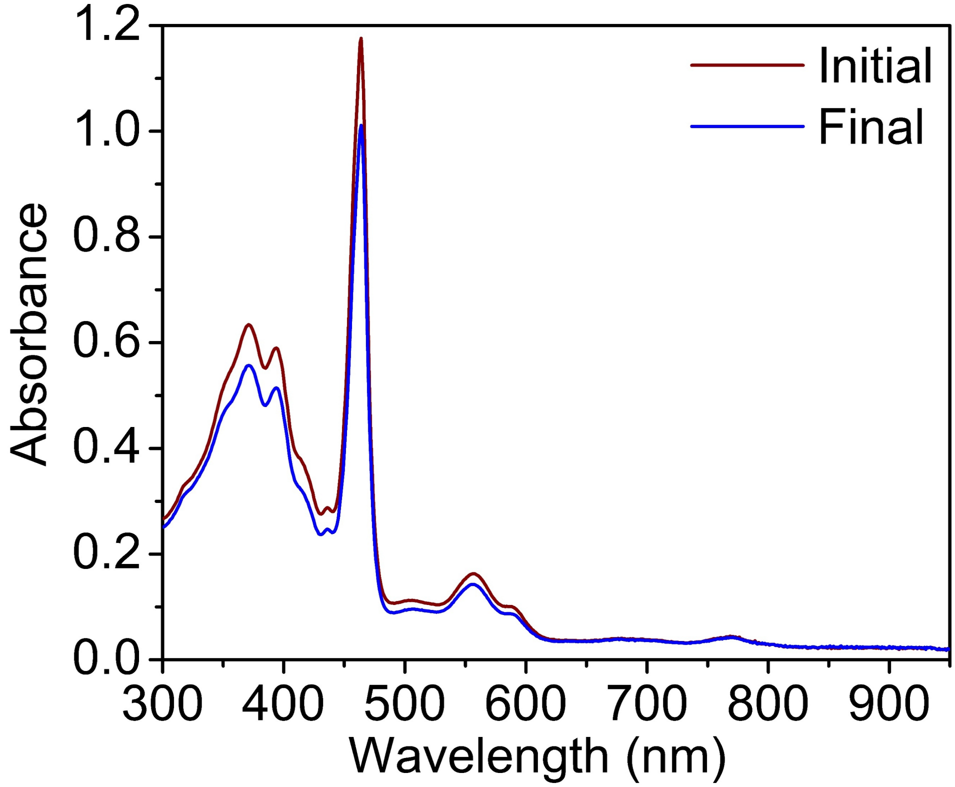


**Figure S21**. UV-vis spectrum of Mn(III)-MPP1 following reaction with $NH_2OH$ as the aminating reagent, showing no significant spectral changes relative to the starting complex.

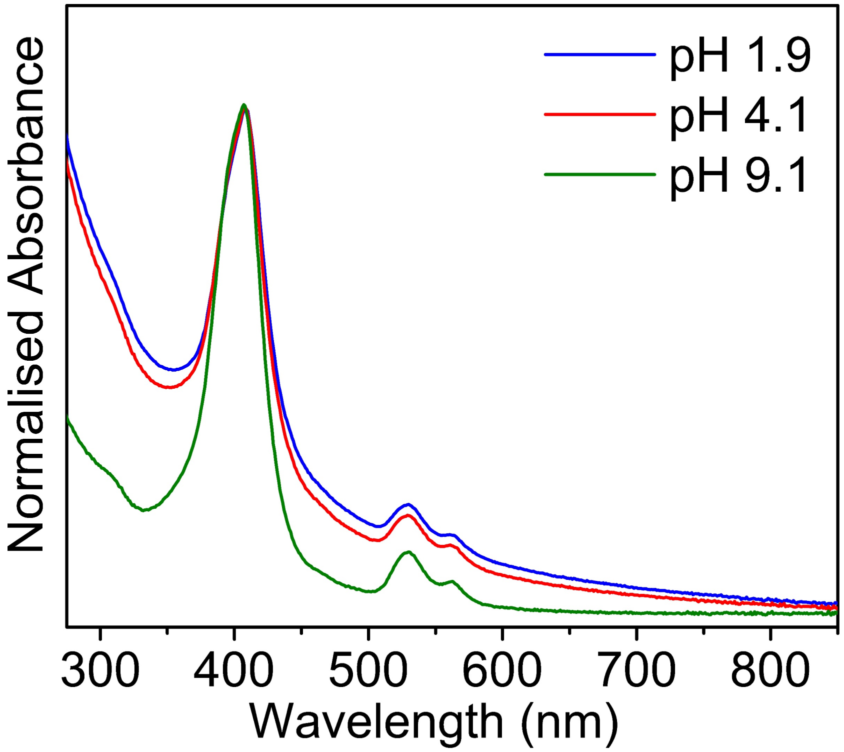


**Figure S22**. UV-vis spectra of Mn(V)(N)MPP1 collected across a range of pH values, demonstrating the stability of the Mn(V)-nitrido species over a wide pH window. The pH of the solution was adjusted by incremental addition of NaOH to universal buffer (40 mM citrate, 40 mM phosphate, 40 mM boric acid, 200 mM NaCl), followed by equilibration of Mn(V)(N)MPP1 at each respective pH prior to spectral acquisition.

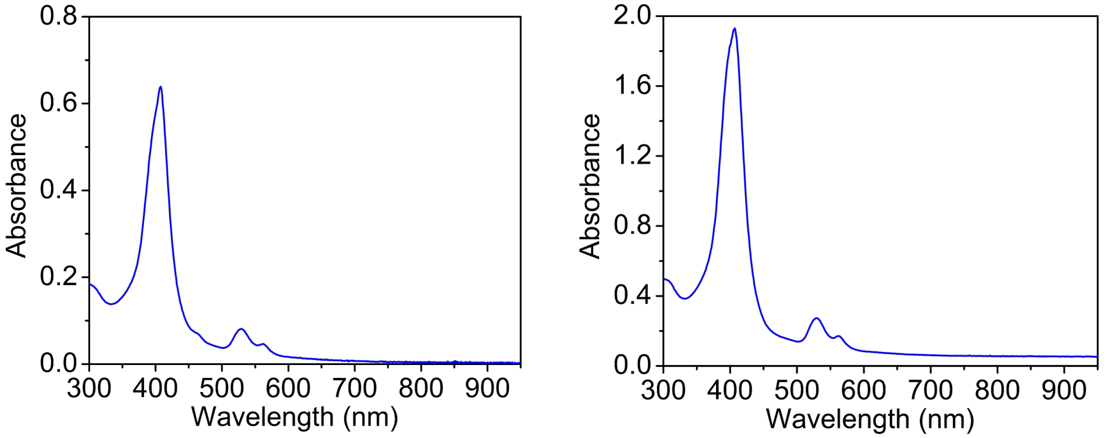


**Figure S23**. UV-vis spectra of the Raman samples of Mn($^{14}$N)-MPP1 (left) and Mn($^{15}$N)-MPP1 (right), recorded before freezing the sample.

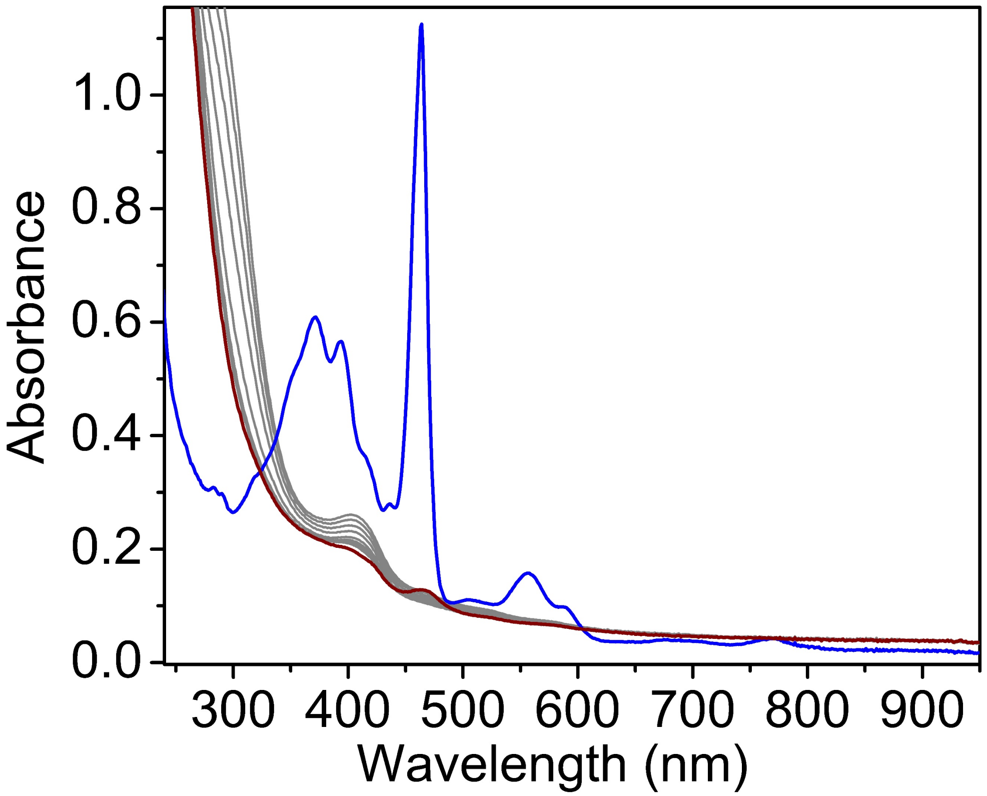


**Figure S24**. UV-vis spectrum of Mn(III)MPP1 upon treatment with excess NaOCl (475 mM, same amount as in catalytic condition) at room temperature. Rapid loss of characteristic absorption features indicates immediate decomposition of the holo protein in the absence of $NH_4OH$.

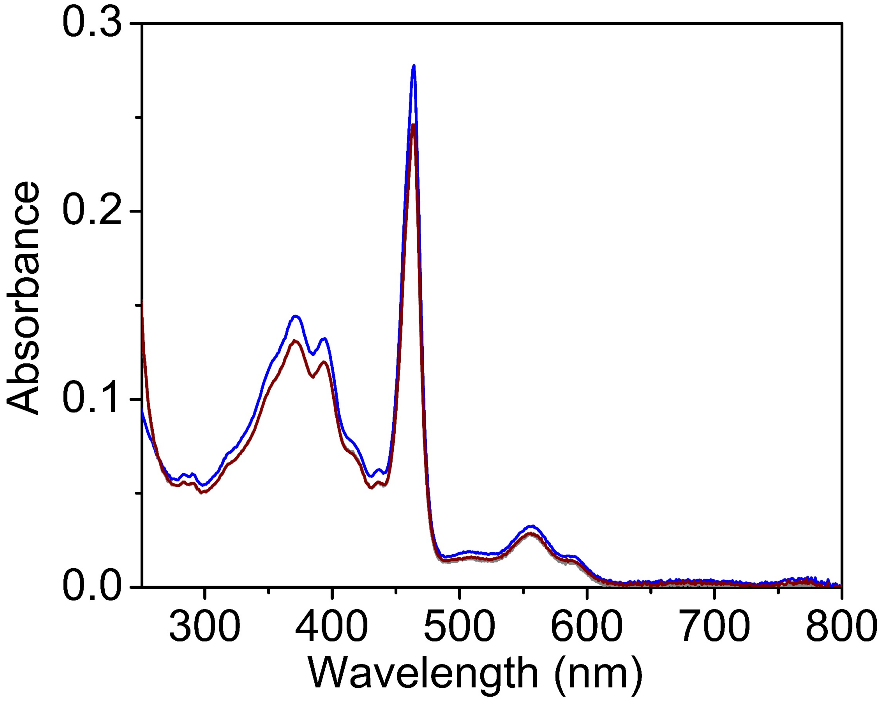


**Figure S25**. UV-vis spectra of Mn(III)MPP1 before (initial, blue) and after (final, red) reaction with *O*-pivaloylhydroxyamine triflic acid at room temperature, showing no reaction of the holo protein under these conditions.

**Reactivity Studies – Calibration Curve for N-Chloro Phenyl Aziridine and GC-MS trace**

Calibration curves of authentic products were used to quantify product formation and determine the yield of reactions involving the proteins. Stock solutions of authentic products (1-16 mM in EtOAc) were prepared by serial dilution and internal standard was added to all the samples. The samples were transferred to 400 μL inserts in 2 mL screw-top vials and analyzed by GC-MS. The calibration curves plot product concentration in mM (y-axis) against the ratio of product peak area over internal standard peak area from GC-MS analysis (x-axis). All data points represent the average of triplicate runs.

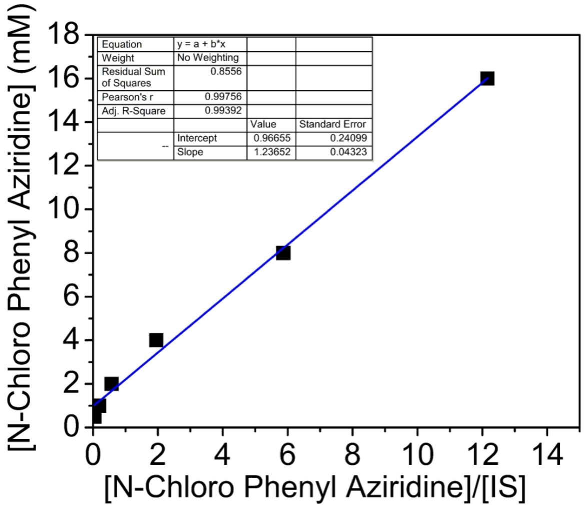


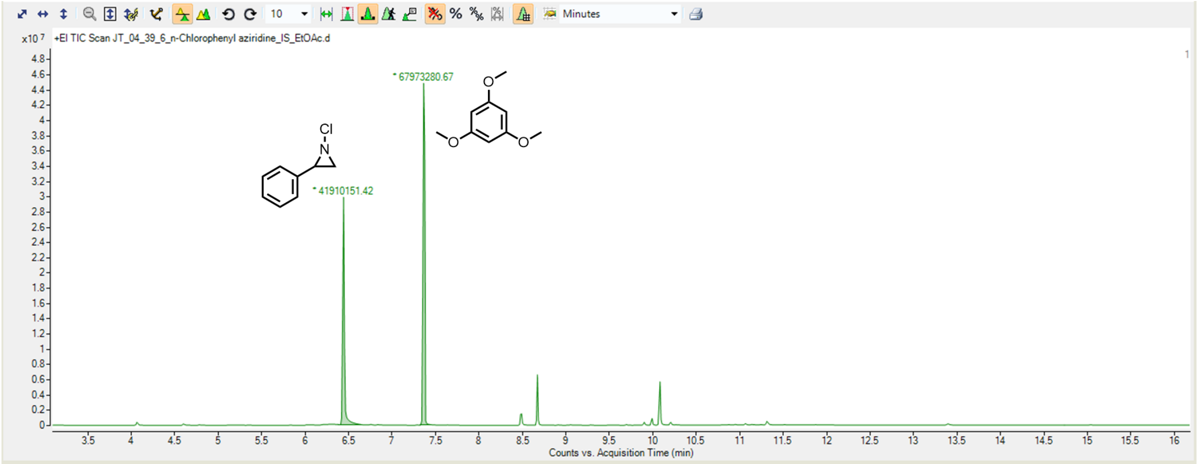


**Figure S26**. GC-MS traces showing the N-chloro-2-phenyl aziridine product (retention time = 6.3 min) and the internal standard (retention time = 7.3 min).

### Reactivity Studies – Calibration Curve for 2-Phenyl Aziridine and GC-MS trace

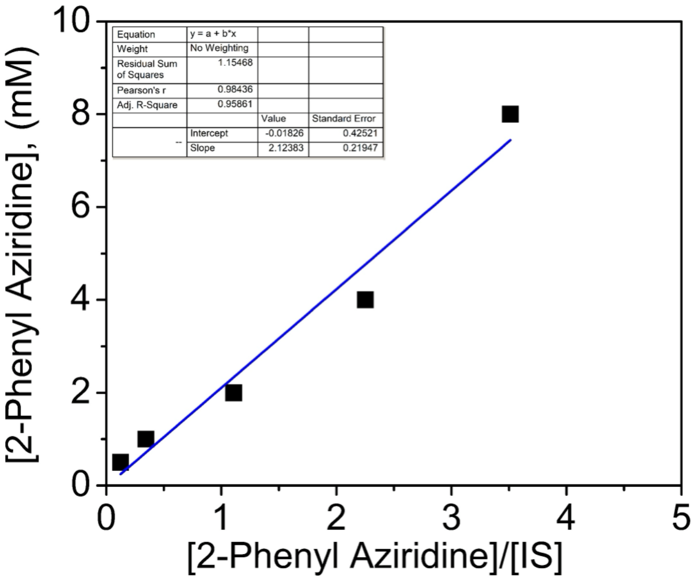


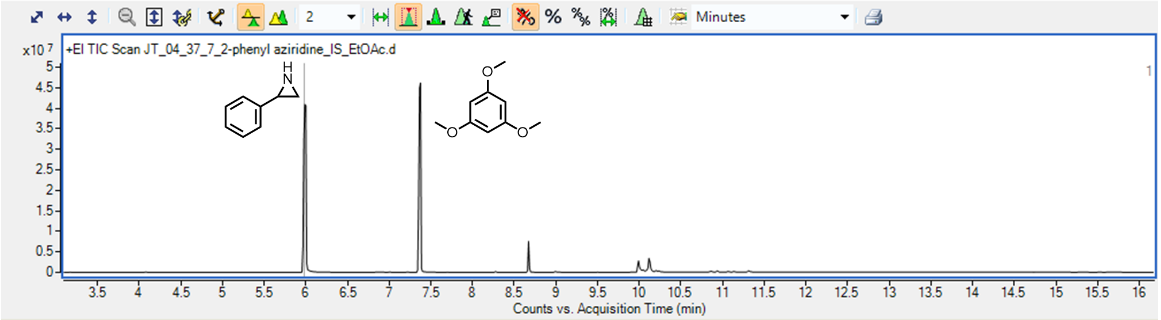


**Figure S27**. GC-MS traces showing 2-phenyl aziridine product (retention time = 5.9 minutes) and internal standard (retention time = 7.3 minutes).

**Reactivity Studies – Calibration Curve for Phenyldichloroethane and GC-MS trace**

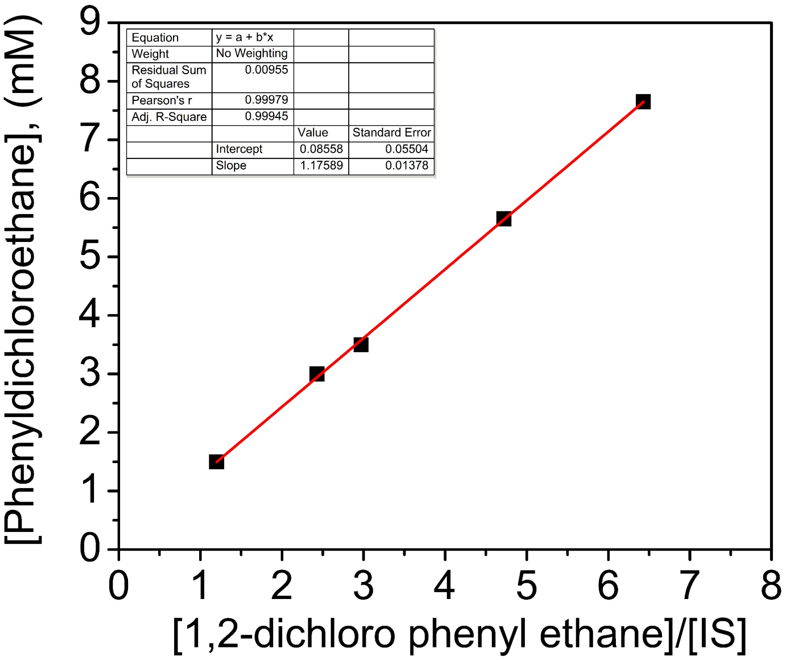


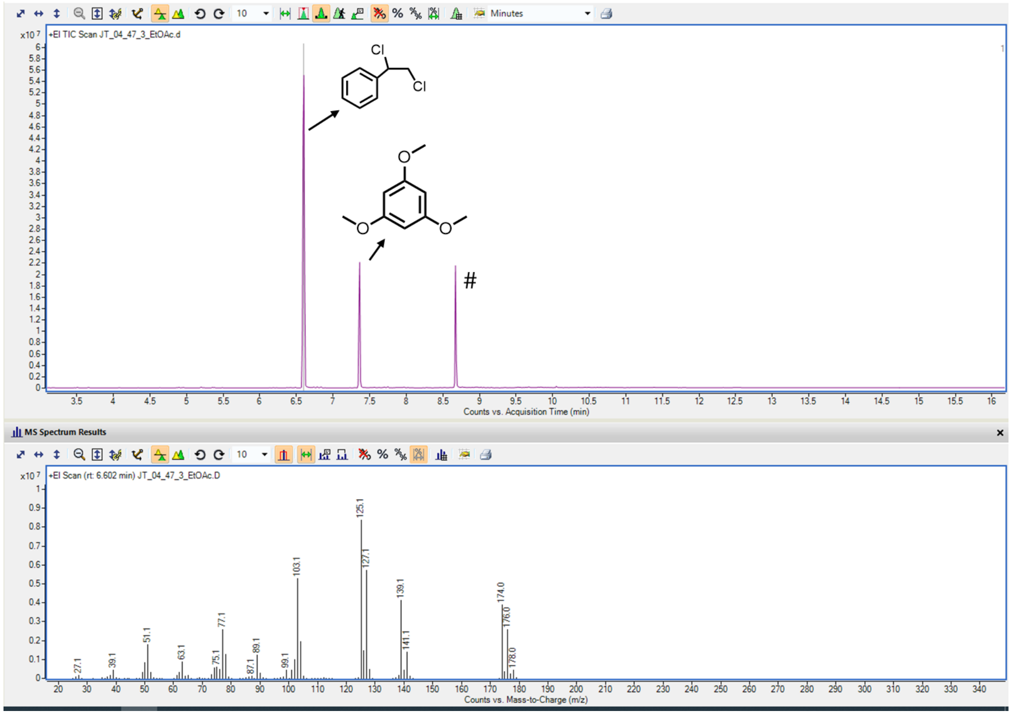


**Figure S28**. GC-MS traces of commercially available Phenyldichloroethane product (retention time = 6.6 minutes) and internal standard (retention time = 7.3 minutes). The (#) corresponds to the impurity in the GC-MS column.

**Table S4. Reactivity studies: effect of catalyst concentration.**

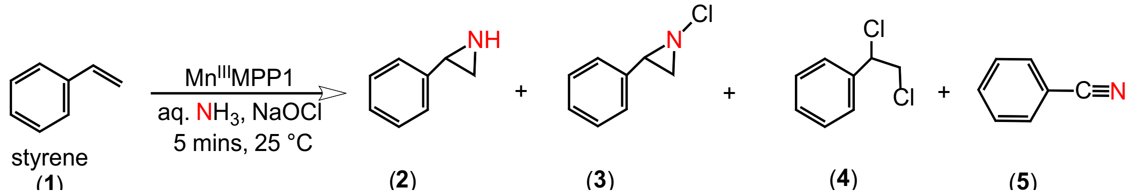


| Catalyst concentration (µM Mn$^{III}$MPP1) | 2 | 3 | 4 | 5 |
|---|---|---|---|---|
| 5 | n.d. | 37 % | 4 | n.d. |
| 25 | n.d. | 50 % | 30 % | n.d. |
| 50 | n.d. | 50 % | 38 % | n.d. |
| 100 | 10 % | 42 % | 10 % | n.d. |
| 200 | 49 % | 35 % | 15 % | n.d. |

**Standard conditions**: Catalyst concentration = 5 - 200 µM, $NH_4OH$ = 740 mM, NaOCl = 475 mM, styrene = 10 mM in MeOH, 1 mL total volume, reaction time = 5 minutes.

n.d. means not determined due to negligible amount.

### GC-MS traces showing formation of products at different catalyst concentrations

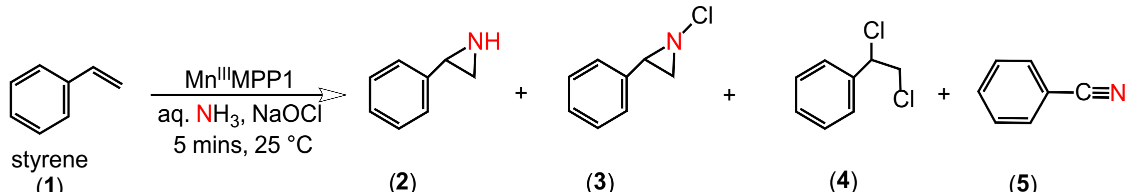


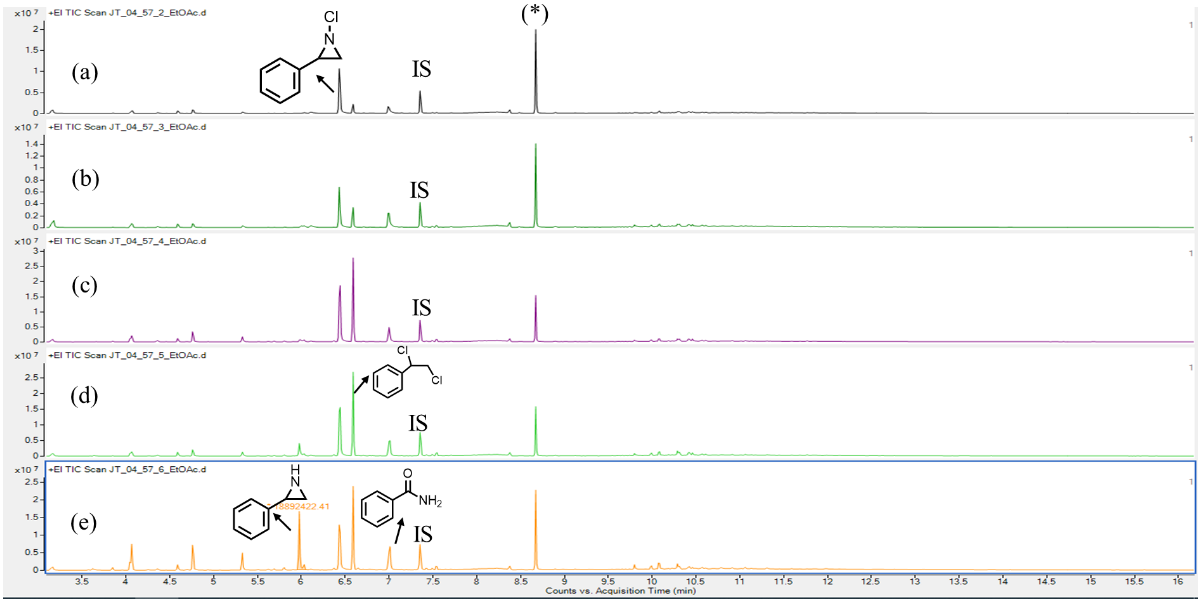


**Figure S29**. GC–MS traces showing product formation at different catalyst concentrations: (a) 5 µM, (b) 25 µM, (c) 50 µM, (d) 100 µM, and (e) 200 µM. Reaction conditions: $NH_4OH$ (740 mM), NaOCl (475 mM), styrene (10 mM) in MeOH (1 mL total volume), reaction time = 5 min. The asterisk (*) denotes an unidentified column-related impurity at a retention time of 8.6 min. IS corresponds to the internal standard, 1,3,5-trimethoxybenzene, observed at a retention time of 7.3 min.

**Table S5. Reactivity Studies: Effect of Different Nitrogen Sources**

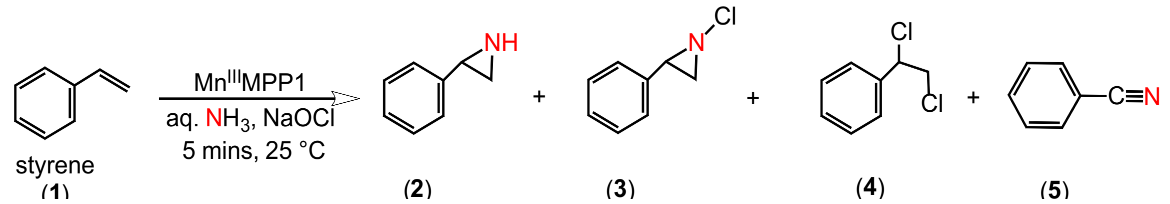


| N sources (740 mM) | 2 | 3 | 4 | 5 |
|---|---|---|---|---|
| $NH_4Cl$ | nd | nd | nd | nd |
| $NH_4OH$ | nd | 50 % | 30 % | nd |
| $NH_2OH$ | nd | nd | nd | nd |
| $NH_2Cl^*$ | nd | nd | nd | nd |
| $NH_4Cl$ + NaOH | nd | 48 % | 27 % | nd |

**Standard conditions**: Catalyst concentration = 25 µM, nitrogen sources = 740 mM, NaOCl = 475 mM, styrene = 10 mM in MeOH, 1 mL total volume, reaction time = 5 minutes. * $NH_2Cl$ = 7.5 mM.

**Table S6. Reactivity Studies with Different Oxidant Sources**

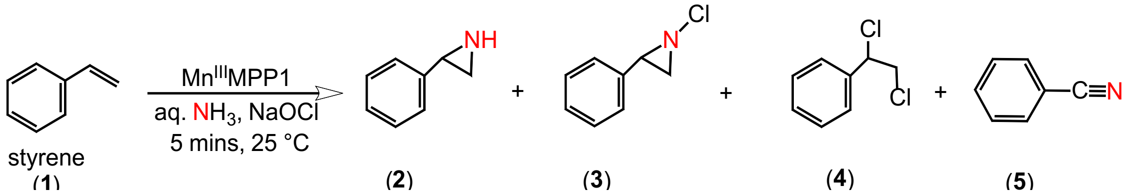


| Oxidant sources (475 mM) | 2 | 3 | 4 | 5 |
|---|---|---|---|---|
| NaOCl | - | 50 % | 30 % | n.d. |
| NaOBr* | Not observed | Not observed | Not observed | Not observed |
| NaIO4 | Not observed | Not observed | Not observed | Not observed |
| $H_2O_2$ | Not observed | Not observed | Not observed | Not observed |

**Standard conditions**: Catalyst concentration = 25 µM, $NH_4OH$= 740 mM, Oxidant sources = 475 mM, styrene = 10 mM in MeOH, 1 mL total volume, reaction time = 5 minutes. * NaOBr = 25 mM

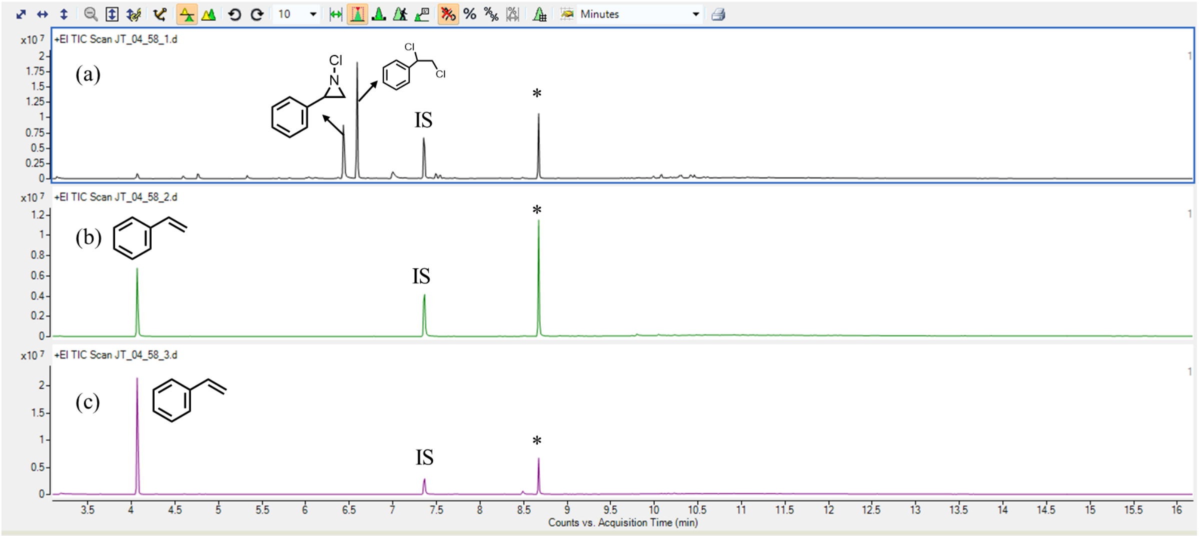


**Figure S30**. GC-MS traces showing the reaction of MnMPP1 and styrene under standard conditions using oxidants (a) NaOCl (b) $NaIO_4$ (c) $H_2O_2$.

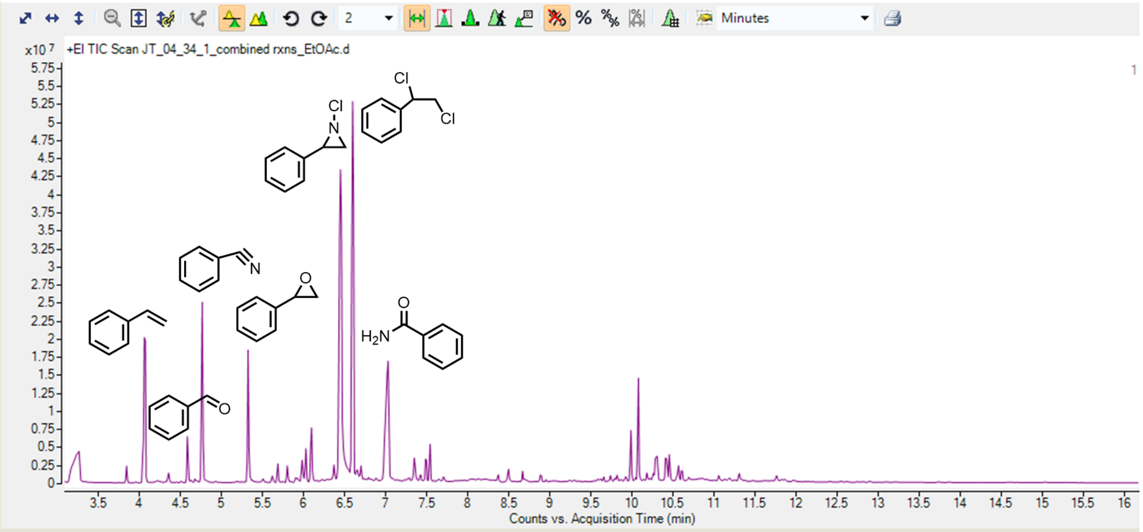


**Figure S31**. GC-MS Analysis of the crude reaction mixture prepared by combining five individual reactions with $Mn^{III}$MPP1 concentration = 25 µM, $NH_4OH$ = 740 mM, NaOCl = 475 mM, styrene = 10 mM in MeOH, 1 mL total volume, reaction time = 5 minutes prior to analysis.

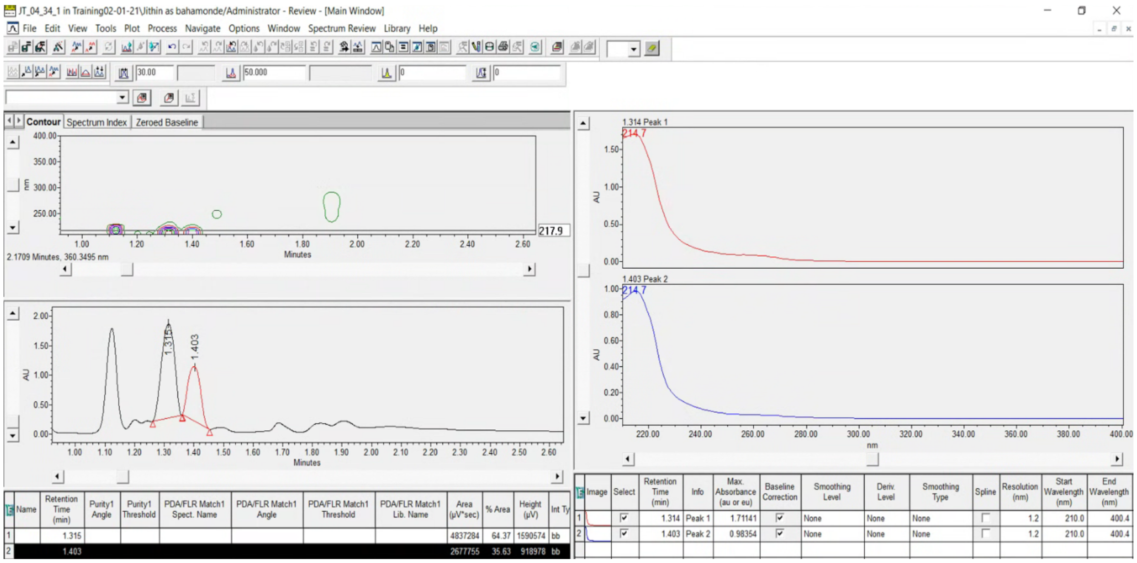


**Figure S32**. UPC Analysis of the above same sample showing enantioselective N-Chloro phenyl aziridine product at a retention time of 1.35 minutes and 1.40 minutes with an ee of 30 %. The peak corresponding to the formation of Phenyldichloroethane can also be observed at a retention time of 1.15 minutes. % ee determined as $(S - R)/(S + R)$.

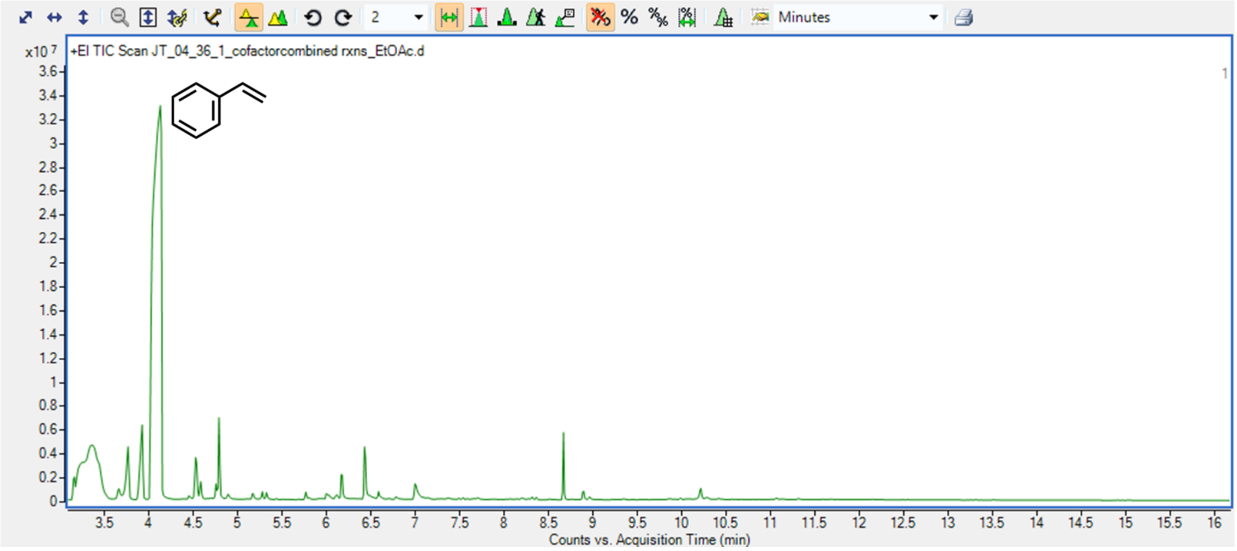


**Figure S33**. GC-MS of the crude reaction mixture prepared by combining five individual reactions with Mn$^{III}$DPP(Cl) cofactor concentration = 25 µM, $NH_4OH$ = 740 mM, NaOCl = 475 mM, styrene = 10 mM in MeOH, 1 mL total volume, reaction time = 5 minutes prior to analysis showing the starting material styrene and negligible amount of product **3** formation.

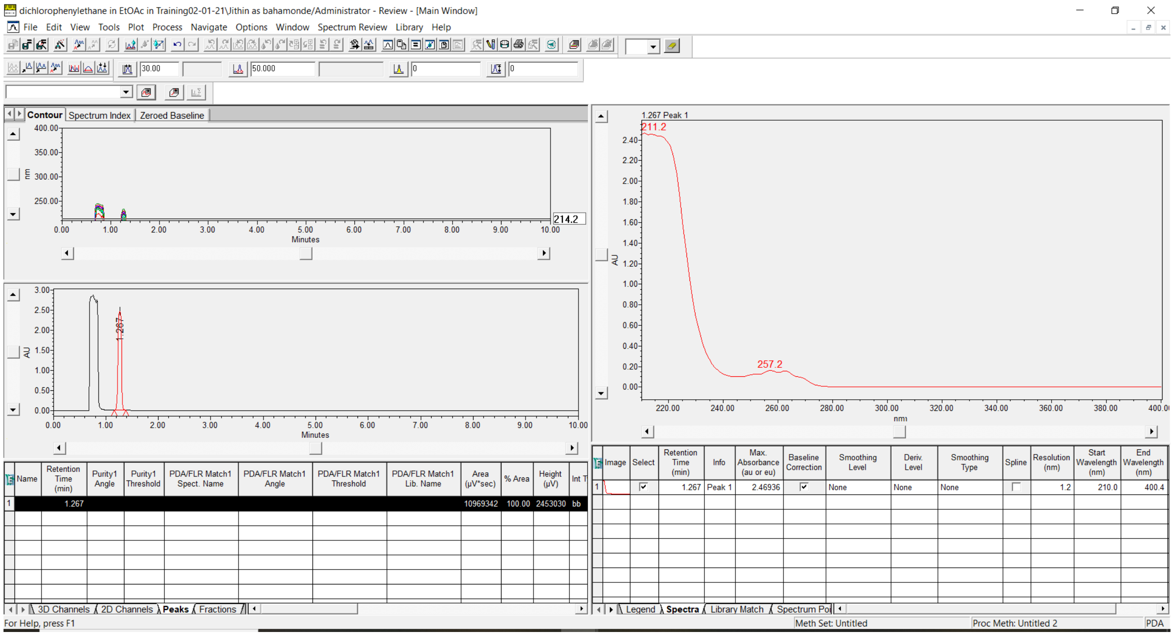

**Figure S34**. UPC analysis of commercially available Phenyldichloroethane with a retention time = 1.20 minutes.

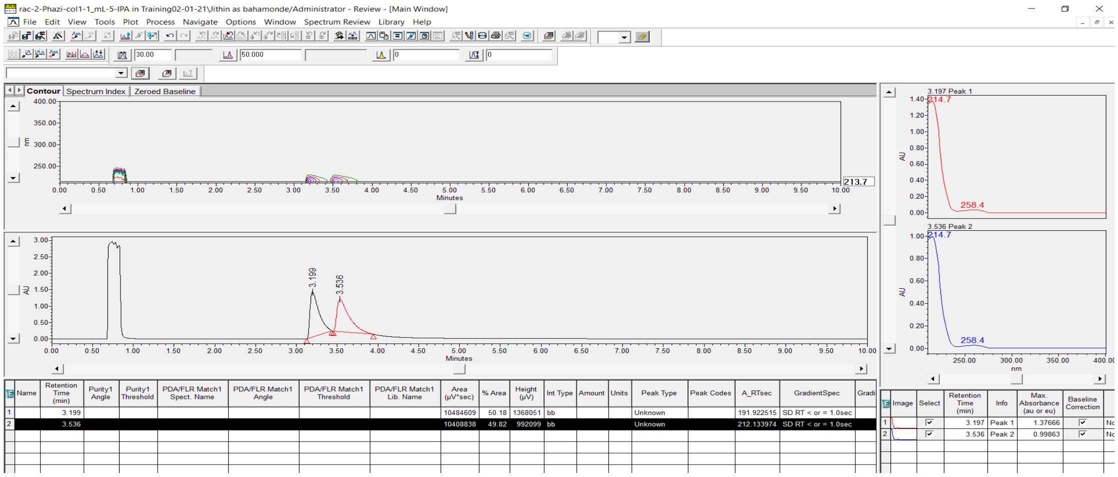

**Figure S35**. UPC analysis of racemic 2- phenyl aziridine product with a retention time = 3.2 and 3.5 minutes.

### Determination of Absolute Configuration

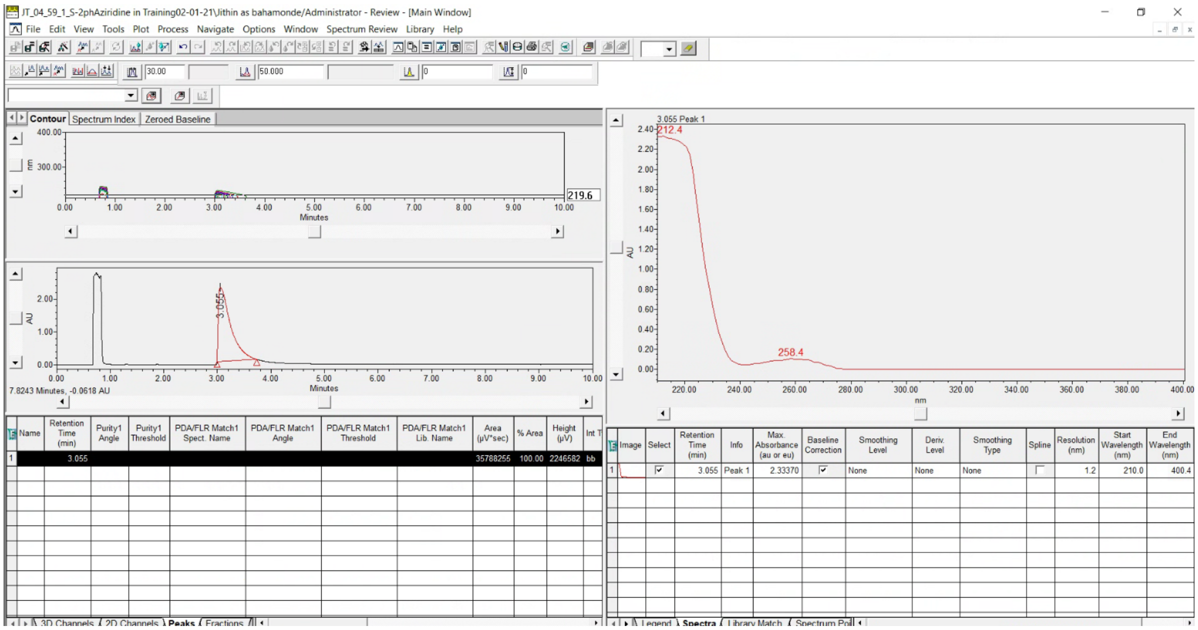

**Figure S36**. UPC analysis of standard (S)- 2-phenyl aziridine

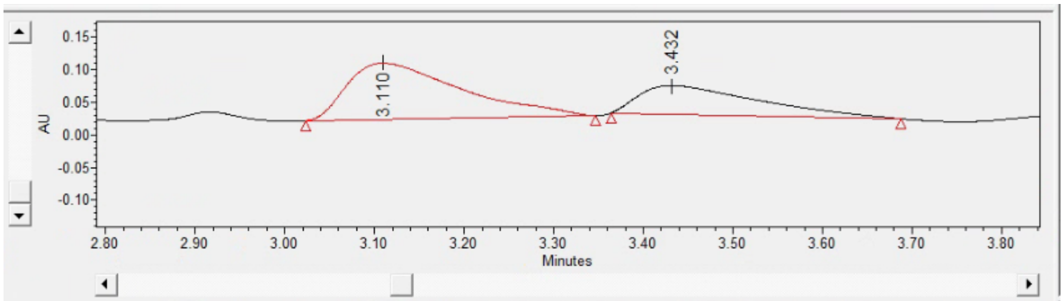


| | Name | Retention Time (min) | Purity1 Angle | Purity1 Threshold | PDA/FLR Match1 Spect. Name | PDA/FLR Match1 Angle | PDA/FLR Match1 Threshold | PDA/FLR Match1 Lib. Name | Area (µV*sec) | % Area | Height (µV) | Int Type |
|---|---|---|---|---|---|---|---|---|---|---|---|---|
| 1 | | 3.110 | | | | | | | 760892 | 65.06 | 85991 | bb |
| 2 | | 3.432 | | | | | | | 408546 | 34.94 | 43750 | bb |

3D Channels | 2D Channels | **Peaks** | Fractions

**Figure S37.** UPC analysis showing enantioselective formation of 2-phenylaziridine product during the reaction of Mn(III)MPP1 (200 µM) with styrene in the presence of $NH_4OH$ and NaOCl after 5 minutes with a preference for S-enantiomer product. Absolute configurations were assigned based on analogy to standard product as shown in **Figure S27**. Enantiomeric ratio of 2-Phenyl Aziridine (S:R) = 65:35. The preference for S-enantiomer of 2-Phenyl aziridine is consistent with that is observed for aziridination using evolved P450 enzymes.[2]

### X-ray Diffraction Data collection

Single crystals suitable for X-ray diffraction were grown by vapor diffusion of hexane into chlorobenzene. A red crystal (plate, approximate dimensions 0.08 × 0.02 × 0.01 $mm^3$) was placed onto the tip of a MiTeGen pin and mounted on a Bruker Venture D8 diffractometer equipped with a PhotonIII detector at 100.00 K. The data collection was carried out using Cu Ka radiation (l = 1.54178 Å, ImS micro-source) with a frame time of 4 seconds and a detector distance of 40 mm. A collection strategy was calculated and complete data to a resolution of 0.81 Å were collected. The frames were integrated with the Bruker SAINT[9] software package using a narrow-frame algorithm to 0.83 Å resolution. Data were corrected for absorption effects using the multi-scan method (SADABS).[10] Please refer to Table S5-S7 for additional crystal and refinement information.

### Structure solution and refinement

The space group P $2_1$/c was determined based on intensity statistics and systematic absences. The structure was solved using the SHELX suite of programs[11-12] and refined using full-matrix least-squares on $F^2$ within the OLEX2 suite.[13] An intrinsic phasing solution was calculated, which provided most non-hydrogen atoms from the E-map. Full-matrix least squares / difference Fourier cycles were performed, which located the remaining non-hydrogen atoms. All non-hydrogen atoms were refined with anisotropic displacement parameters. The hydrogen atoms were placed in ideal positions and refined as riding atoms with relative isotropic displacement parameters. Only half of the molecule belongs to the asymmetric unit. The other half is generated by symmetry and the Mn atom and nitride ligand are equally disordered over two positions above and below the porphyrin plane, similarly to structures reported for other Mn-porphyrin complexes.[14] The final full matrix least squares refinement converged to R1 = 0.0480 and wR2 = 0.1042 ($F^2$, all data). The goodness-of-fit was 1.112. Based on the final model, the calculated density was 1.475 $g/cm^3$ and F(000), 544 $e^-$.

**Table S5. Crystal data and structure refinement for Mn(V)(N) DPP.**

| | | |
|---|---|---|
| Empirical formula | C32 H20 Mn N5 | |
| Formula weight | 529.47 | |
| Crystal color, shape, size | red plate, 0.08 × 0.02 × 0.01 $mm^3$ | |
| Temperature | 100.00 K | |
| Wavelength | 1.54178 Å | |
| Crystal system, space group | Monoclinic, P $2_1$/c | |
| Unit cell dimensions | a = 6.7876(3) Å | a = 90°. |
| | b = 18.9334(7) Å | b = 105.804(3)°. |
| | c = 9.6393(4) Å | g = 90°. |
| Volume | 1191.94(9) $Å^3$ | |
| Z | 2 | |
| Density (calculated) | 1.475 $g/cm^3$ | |
| Absorption coefficient | 4.758 $mm^{-1}$ | |
| F(000) | 544 | |
| **Data collection** | | |
| Diffractometer | Bruker D8 Venture | |
| Theta range for data collection | 4.671 to 68.335°. | |
| Index ranges | -8<=h<=8, -22<=k<=22, -11<=l<=11 | |
| Reflections collected | 22250 | |
| Independent reflections | 2175 [Rint = 0.0654] | |
| Observed Reflections | 1868 | |
| Completeness to theta = 67.679° | 99.9 % | |
| **Solution and Refinement** | | |
| Absorption correction | Semi-empirical from equivalents | |
| Max. and min. transmission | 0.7531 and 0.6258 | |
| Solution | Intrinsic methods | |
| Refinement method | Full-matrix least-squares on $F^2$ | |
| Weighting scheme | w = $[s^2Fo^2 + AP^2 + BP]^{-1}$, with | |
| | P = $(Fo^2 + 2 Fc^2)/3$, A = 0.005, B = 2.67 | |
| Data / restraints / parameters | 2175 / 0 / 181 | |
| Goodness-of-fit on $F^2$ | 1.112 | |
| Final R indices [I>2s(I)] | R1 = 0.0480, wR2 = 0.0992 | |
| R indices (all data) | R1 = 0.0576, wR2 = 0.1042 | |
| Largest diff. peak and hole | 0.272 and -0.390 e.$Å^{-3}$ | |

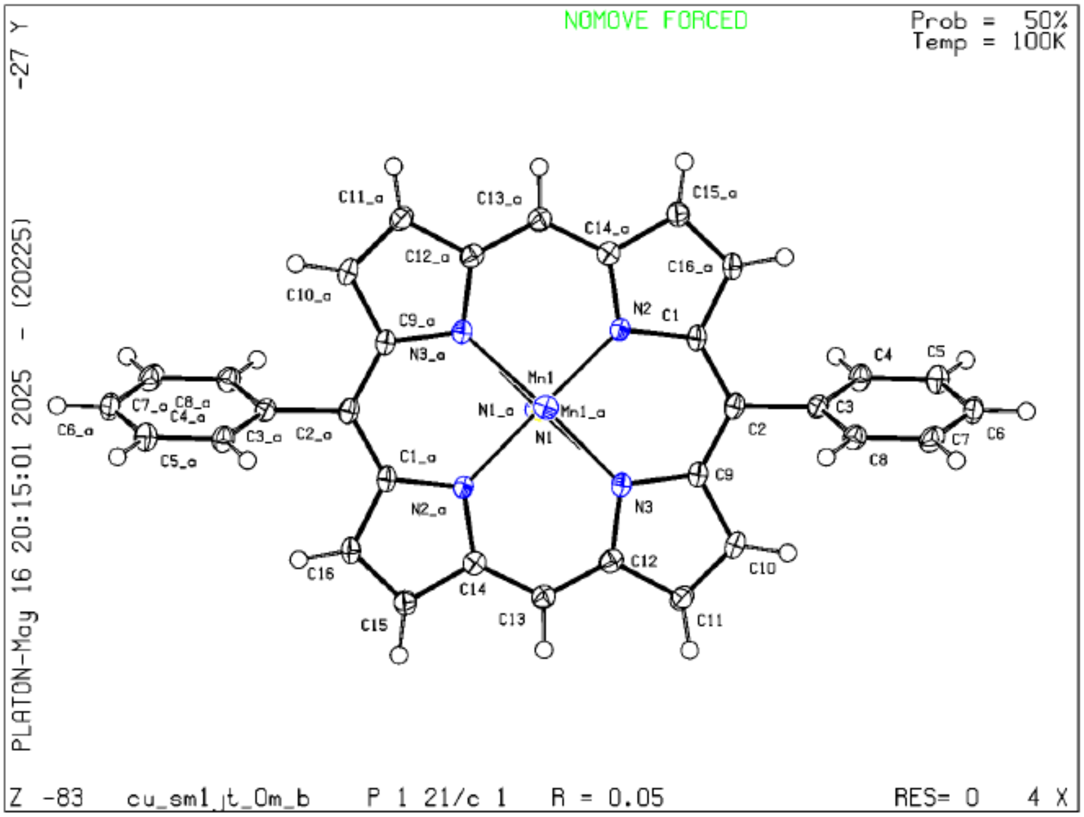
NOMOVE FORCED
Prob = 50%
Temp = 100K
-27 Y
PLATON-May 16 20:15:01 2025 - (20225)
Z -83 cu_sm1_jt_0m_b P 1 21/c 1 R = 0.05 RES= 0 4 X
C11_a
C13_a
C15_a
C14_a
C12_a
C16_a
C10_a
N2
C1
C9_a
N3_a
C4
C5
Mn1
C7_a C8_a
C4_a
N1_a
Mn1_a
C3
C6_a
C3_a
C2_a
N1
C2
C6
C7
C8
C5_a
C1_a
C9
N2_a
N3
C16
C14
C12
C10
C15
C13
C11

**Table S6. Bond Parameters of Mn(V)N-DPP**

| Atom1 | Atom2 | Length |
|---|---|---|
| N2 | C1 | 1.385(3) |
| N2 | Mn1 | 1.972(2) |
| N2 | C14 | 1.378(3) |
| N2 | Mn1 | 2.088(2) |
| N3 | C9 | 1.386(3) |
| N3 | C12 | 1.380(3) |
| N3 | Mn1 | 2.058(2) |
| N3 | Mn1 | 1.983(2) |
| C1 | C2 | 1.396(4) |
| C1 | C16 | 1.435(4) |
| C2 | C3 | 1.497(4) |
| C2 | C9 | 1.391(4) |
| C3 | C4 | 1.392(4) |
| C3 | C8 | 1.395(4) |
| C4 | C5 | 1.390(4) |
| C5 | C6 | 1.386(4) |
| C6 | C7 | 1.387(4) |
| C7 | C8 | 1.389(4) |
| N3 | C12 | 1.380(3) |
| N3 | Mn1 | 2.058(2) |
| C1 | C2 | 1.396(4) |
| C2 | C3 | 1.497(4) |
| C2 | C9 | 1.391(4) |
| C3 | C4 | 1.392(4) |
| C3 | C8 | 1.395(4) |
| C4 | C5 | 1.390(4) |
| C5 | C6 | 1.386(4) |
| C6 | C7 | 1.387(4) |
| C7 | C8 | 1.389(4) |
| C9 | C10 | 1.437(4) |
| C10 | C11 | 1.352(4) |
| C11 | C12 | 1.434(4) |
| C12 | C13 | 1.379(4) |
| C13 | C14 | 1.382(4) |
| C14 | C15 | 1.430(4) |
| C15 | C16 | 1.356(4) |
| Mn1 | N1 | 1.534(4) |
| C9 | C10 | 1.437(4) |
| C10 | C11 | 1.352(4) |
| C11 | C12 | 1.434(4) |
| C12 | C13 | 1.379(4) |
| C13 | C14 | 1.382(4) |
| C14 | C15 | 1.430(4) |
| C14 | N2 | 1.378(3) |
| C15 | C16 | 1.356(4) |
| C16 | C1 | 1.435(4) |
| Mn1 | N1 | 1.534(4) |
| Mn1 | N2 | 2.088(2) |
| Mn1 | N3 | 1.983(2) |
| Mn1 | Mn1 | 0.778(1) |
| Mn1 | N1 | 2.305(4) |
| N1 | Mn1 | 2.305(4) |
| N2 | C1 | 1.385(3) |
| N2 | Mn1 | 1.972(2) |
| N3 | C9 | 1.386(3) |

**Table S7. Bond Angles of Mn(V)(N)-DPP**

| Atom1 | Atom2 | Atom3 | Angle |
|---|---|---|---|
| C1 | N2 | Mn1 | 128.7(2) |
| C1 | N2 | C14 | 104.8(2) |
| C1 | N2 | Mn1 | 126.9(2) |
| Mn1 | N2 | C14 | 124.4(2) |
| Mn1 | N2 | Mn1 | 21.84(4) |
| C14 | N2 | Mn1 | 127.2(2) |
| C9 | N3 | C12 | 105.1(2) |
| C9 | N3 | Mn1 | 125.6(2) |
| C9 | N3 | Mn1 | 129.6(2) |
| C12 | N3 | Mn1 | 128.0(2) |
| C12 | N3 | Mn1 | 123.3(2) |
| Mn1 | N3 | Mn1 | 22.09(4) |
| N2 | C1 | C2 | 125.2(2) |
| N2 | C1 | C16 | 110.2(2) |
| C2 | C1 | C16 | 124.5(3) |
| C1 | C2 | C3 | 119.2(2) |
| C1 | C2 | C9 | 123.0(3) |
| C3 | C2 | C9 | 117.9(2) |
| C2 | C3 | C4 | 121.3(2) |
| C2 | C3 | C8 | 119.9(2) |
| C4 | C3 | C8 | 118.8(3) |
| C3 | C4 | H4 | 119.8 |
| C3 | C4 | C5 | 120.5(3) |
| H4 | C4 | C5 | 119.8 |
| C4 | C5 | H5 | 119.9 |
| C4 | C5 | C6 | 120.3(3) |
| H5 | C5 | C6 | 119.9 |
| C5 | C6 | H6 | 120.1 |
| C5 | C6 | C7 | 119.8(3) |
| H6 | C6 | C7 | 120.1 |
| C6 | C7 | H7 | 120.1 |
| C6 | C7 | C8 | 119.8(3) |
| H7 | C7 | C8 | 120.1 |
| C3 | C8 | C7 | 120.8(3) |
| C3 | C8 | H8 | 119.6 |
| C7 | C8 | H8 | 119.6 |
| N3 | C9 | C2 | 125.8(2) |
| N3 | C9 | C10 | 110.1(2) |
| C2 | C9 | C10 | 124.1(3) |
| C9 | C10 | H10 | 126.3 |
| C9 | C10 | C11 | 107.3(2) |
| H10 | C10 | C11 | 126.4 |
| C10 | C11 | H11 | 126.5 |
| C10 | C11 | C12 | 106.9(2) |
| H11 | C11 | C12 | 126.5 |
| N3 | C12 | C11 | 110.6(2) |
| N3 | C12 | C13 | 125.3(2) |
| C11 | C12 | C13 | 124.1(3) |
| C12 | C13 | H13 | 117.2 |
| C12 | C13 | C14 | 125.7(3) |
| H13 | C13 | C14 | 117.2 |
| C13 | C14 | C15 | 124.0(3) |
| C13 | C14 | N2 | 124.9(2) |
| C15 | C14 | N2 | 111.2(2) |
| C14 | C15 | H15 | 126.7 |
| C14 | C15 | C16 | 106.5(2) |
| H15 | C15 | C16 | 126.7 |
| C15 | C16 | H16 | 126.3 |
| C15 | C16 | C1 | 107.3(2) |
| H16 | C16 | C1 | 126.4 |
| N2 | Mn1 | N3 | 87.46(9) |
| N2 | Mn1 | N1 | 100.0(2) |
| N2 | Mn1 | N2 | 158.2(1) |
| N2 | Mn1 | N3 | 91.64(9) |
| N2 | Mn1 | Mn1 | 87.5(1) |
| N2 | Mn1 | N1 | 82.5(1) |
| N3 | Mn1 | N1 | 101.5(2) |
| N3 | Mn1 | N2 | 86.34(9) |
| N3 | Mn1 | N3 | 157.9(1) |
| N3 | Mn1 | Mn1 | 73.5(1) |
| N3 | Mn1 | N1 | 76.8(1) |
| N1 | Mn1 | N2 | 101.7(2) |
| N1 | Mn1 | N3 | 100.4(2) |
| N1 | Mn1 | Mn1 | 170.9(2) |
| N1 | Mn1 | N1 | 176.9(2) |
| N2 | Mn1 | N3 | 86.36(9) |
| N2 | Mn1 | Mn1 | 70.7(1) |

| | | | |
|---|---|---|---|
| N2 | Mn1 | N1 | 75.7(1) |
| N3 | Mn1 | Mn1 | 84.4(1) |
| N3 | Mn1 | N1 | 81.2(1) |
| Mn1 | Mn1 | N1 | 6.0(1) |
| Mn1 | N1 | Mn1 | 3.06(7) |
| C14 | N2 | Mn1 | 127.2(2) |
| C14 | N2 | C1 | 104.8(2) |
| C14 | N2 | Mn1 | 124.4(2) |
| Mn1 | N2 | C1 | 126.9(2) |
| Mn1 | N2 | Mn1 | 21.84(4) |
| C1 | N2 | Mn1 | 128.7(2) |
| Mn1 | N3 | C9 | 129.6(2) |
| Mn1 | N3 | C12 | 123.3(2) |
| Mn1 | N3 | Mn1 | 22.09(4) |
| C9 | N3 | C12 | 105.1(2) |
| C9 | N3 | Mn1 | 125.6(2) |
| C12 | N3 | Mn1 | 128.0(2) |
| C16 | C1 | N2 | 110.2(2) |
| C16 | C1 | C2 | 124.5(3) |
| N2 | C1 | C2 | 125.2(2) |
| C1 | C2 | C3 | 119.2(2) |
| C1 | C2 | C9 | 123.0(3) |
| C3 | C2 | C9 | 117.9(2) |
| C2 | C3 | C4 | 121.3(2) |
| C2 | C3 | C8 | 119.9(2) |
| C4 | C3 | C8 | 118.8(3) |
| C3 | C4 | H4 | 119.8 |
| C3 | C4 | C5 | 120.5(3) |
| H4 | C4 | C5 | 119.8 |
| C4 | C5 | H5 | 119.9 |
| C4 | C5 | C6 | 120.3(3) |
| H5 | C5 | C6 | 119.9 |
| C5 | C6 | H6 | 120.1 |
| C5 | C6 | C7 | 119.8(3) |
| H6 | C6 | C7 | 120.1 |
| C6 | C7 | H7 | 120.1 |
| C6 | C7 | C8 | 119.8(3) |
| H7 | C7 | C8 | 120.1 |
| C3 | C8 | C7 | 120.8(3) |
| C3 | C8 | H8 | 119.6 |
| C7 | C8 | H8 | 119.6 |
| N3 | C9 | C2 | 125.8(2) |
| N3 | C9 | C10 | 110.1(2) |
| C2 | C9 | C10 | 124.1(3) |
| C9 | C10 | H10 | 126.3 |
| C9 | C10 | C11 | 107.3(2) |
| H10 | C10 | C11 | 126.4 |
| C10 | C11 | H11 | 126.5 |
| C10 | C11 | C12 | 106.9(2) |
| H11 | C11 | C12 | 126.5 |
| N3 | C12 | C11 | 110.6(2) |
| N3 | C12 | C13 | 125.3(2) |
| C11 | C12 | C13 | 124.1(3) |
| C12 | C13 | H13 | 117.2 |
| C12 | C13 | C14 | 125.7(3) |
| H13 | C13 | C14 | 117.2 |
| N2 | C14 | C13 | 124.9(2) |
| N2 | C14 | C15 | 111.2(2) |
| C13 | C14 | C15 | 124.0(3) |
| C14 | C15 | H15 | 126.7 |
| C14 | C15 | C16 | 106.5(2) |
| H15 | C15 | C16 | 126.7 |
| C1 | C16 | C15 | 107.3(2) |
| C1 | C16 | H16 | 126.4 |
| C15 | C16 | H16 | 126.3 |
| N2 | Mn1 | N3 | 86.36(9) |
| N2 | Mn1 | Mn1 | 70.7(1) |
| N2 | Mn1 | N1 | 75.7(1) |
| N2 | Mn1 | N2 | 158.2(1) |
| N2 | Mn1 | N3 | 86.34(9) |
| N2 | Mn1 | N1 | 101.7(2) |
| N3 | Mn1 | Mn1 | 84.4(1) |
| N3 | Mn1 | N1 | 81.2(1) |
| N3 | Mn1 | N2 | 91.64(9) |
| N3 | Mn1 | N3 | 157.9(1) |
| N3 | Mn1 | N1 | 100.4(2) |
| Mn1 | Mn1 | N1 | 6.0(1) |
| Mn1 | Mn1 | N2 | 87.5(1) |
| Mn1 | Mn1 | N3 | 73.5(1) |
| Mn1 | Mn1 | N1 | 170.9(2) |
| N1 | Mn1 | N2 | 82.5(1) |
| N1 | Mn1 | N3 | 76.8(1) |

| | | | |
|---|---|---|---|
| N1 | Mn1 | N1 | 176.9(2) |
| N2 | Mn1 | N3 | 87.46(9) |
| N2 | Mn1 | N1 | 100.0(2) |
| N3 | Mn1 | N1 | 101.5(2) |
| Mn1 | N1 | Mn1 | 3.06(7) |

### References


1. Stoll, S.; Schweiger, A., EasySpin, a comprehensive software package for spectral simulation and analysis in EPR. *J. Magn. Reson.* **2006,** *178* (1), 42-55.
2. Farwell, C. C.; Zhang, R. K.; McIntosh, J. A.; Hyster, T. K.; Arnold, F. H., Enantioselective Enzyme-Catalyzed Aziridination Enabled by Active-Site Evolution of a Cytochrome P450. *ACS Cent. Sci.* **2015,** *1* (2), 89-93.
3. Polizzi, N. F.; Eibling, M. J.; Perez-Aguilar, J. M.; Rawson, J.; Lanci, C. J.; Fry, H. C.; Beratan, D. N.; Saven, J. G.; Therien, M. J., Photoinduced Electron Transfer Elicits a Change in the Static Dielectric Constant of a de Novo Designed Protein. *J. Am. Chem. Soc.* **2016,** *138* (7), 2130-3.
4. Hynes, J., Jr.; Doubleday, W. W.; Dyckman, A. J.; Godfrey, J. D., Jr.; Grosso, J. A.; Kiau, S.; Leftheris, K., N-Amination of pyrrole and indole heterocycles with monochloramine ($NH_2Cl$). *J. Org. Chem.* **2004,** *69* (4), 1368-71.
5. Stone, K.; Hua, J.; Choudhry, H., Manganese-Substituted Myoglobin: Characterization and Reactivity of an Oxidizing Intermediate towards a Weak C-H Bond. *Inorganics* **2015,** *3* (2), 219-229.
6. Mann, S. I.; Nayak, A.; Gassner, G. T.; Therien, M. J.; DeGrado, W. F., De Novo Design, Solution Characterization, and Crystallographic Structure of an Abiological Mn-Porphyrin-Binding Protein Capable of Stabilizing a Mn(V) Species. *J. Am. Chem. Soc.* **2021,** *143* (1), 252-259.
7. Hill, C. L.; Hollander, F. J., Structural characterization of a complex of Manganese(V) nitrido[tetrakis(p-methoxyphenyl)porphinato] manganese(V). *J. Am. Chem. Soc.* **1982,** *104* (25), 7318-7319.
8. Panetti, G. B.; Kim, J.; Myong, M. S.; Bird, M. J.; Scholes, G. D.; Chirik, P. J., Photodriven Ammonia Synthesis from Manganese Nitrides: Photophysics and Mechanistic Investigations. *J. Am. Chem. Soc.* **2024,** *146* (40), 27610-27621.
9. *SAINT*, V8.30A; Bruker Analytical X-Ray Systems: Madison, WI, 2012.
10. *SADABS*, 2.03; Bruker Analytical X-Ray Systems: Madison, WI, 2016.
11. Sheldrick, G. M., Crystal structure refinement with SHELXL. *Acta Cryst.* **2015,** *A71* (Pt 1), 3-8.
12. Sheldrick, G. M., A short history of SHELX. *Acta Cryst.* **2008,** *A64* (Pt 1), 112-22.
13. Dolomanov, O. V.; Bourhis, L. J.; Gildea, R. J.; Howard, J. A. K.; Puschmann, H., OLEX2: a complete structure solution, refinement and analysis program. *J. Appl. Crystallogr.* **2009,** *42* (2), 339-341.
14. Shields, M. R.; Guzei, I. A.; Goll, J. G., Crystal structure of nitrido[5,10,15,20-tetra-kis(4-methylphenyl)-porphyrinato]-manganese(V). *Acta Cryst.* **2014,** *E70* (Pt 10), 242-245.
